# Comparing DNA replication programs reveals large timing shifts at centromeres of endocycling cells in maize roots

**DOI:** 10.1101/2020.01.24.917914

**Authors:** Emily E. Wear, Jawon Song, Gregory J. Zynda, Leigh Mickelson-Young, Chantal LeBlanc, Tae-Jin Lee, David O. Deppong, George C. Allen, Robert A. Martienssen, Matthew W. Vaughn, Linda Hanley-Bowdoin, William F. Thompson

## Abstract

Plant cells undergo two types of cell cycles – the mitotic cycle in which DNA replication is coupled to mitosis, and the endocycle in which DNA replication occurs in the absence of cell division. To investigate DNA replication programs in these two types of cell cycles, we pulse labeled intact root tips of maize (*Zea mays*) with 5-ethynyl-2’-deoxyuridine (EdU) and used flow sorting of nuclei to examine DNA replication timing (RT) during the transition from a mitotic cycle to an endocycle. Here, we compare sequence-based RT profiles and found that most regions of the maize genome replicate at the same time during S phase in mitotic and endocycling cells, despite the need to replicate twice as much DNA in the endocycle. However, regions collectively corresponding to 2% of the genome displayed significant changes in timing between the two types of cell cycles. The majority of these regions are small, with a median size of 135 kb, and shift to a later RT in the endocycle. However, we found larger regions that shifted RT in centromeres of seven of the ten maize chromosomes. These regions covered the majority of the previously defined functional centromere in each case, which are ∼1–2 Mb in size in the reference genome. They replicate mainly during mid S phase in mitotic cells, but primarily in late S phase of the endocycle. Strikingly, the immediately adjacent pericentromere sequences are primarily late replicating in both cell cycles. Analysis of CENH3 enrichment levels in nuclei of different ploidies suggested that there is only a partial replacement of CENH3 nucleosomes after endocycle replication is complete. The shift to later replication of centromeres and reduced CENH3 enrichment after endocycle replication is consistent with the hypothesis that centromeres are being inactivated as their function is no longer needed.

**AUTHOR SUMMARY:** In traditional cell division, or mitosis, a cell’s genetic material is duplicated and then split between two daughter cells. In contrast, in some specialized cell types, the DNA is duplicated a second time without an intervening division step, resulting in cells that carry twice as much DNA – a phenomenon called an endocycle, which is common during plant development. At each step, DNA replication follows an ordered program, in which highly compacted DNA is unraveled and replicated in sections at different times during the synthesis (S) phase. In plants, it is unclear whether traditional and endocycle programs are the same. Using root tips of maize, we found a small portion of the genome whose replication in the endocycle is shifted in time, usually to later in S phase. Some of these regions are scattered around the genome, and mostly coincide with active genes. However, the most prominent shifts occur in centromeres. This location is noteworthy because centromeres orchestrate the process of separating duplicated chromosomes into daughter cells, a function that is not needed in the endocycle. Our observation that centromeres replicate later in the endocycle suggests there is an important link between the time of replication and the function of centromeres.

## INTRODUCTION

Developmentally programmed DNA replication without nuclear breakdown, chromosome condensation or cell division, a phenomenon known as endoreduplication or endocycling, occurs in a wide variety of plants and animals [1–3]. In plants, endoreduplication is a systemic feature [4] and often an important step in the development of tissues and organs such as fruit, endosperm, leaf epidermal cells, and trichomes [5]. Initiation of endocycling is frequently associated with a transition from cell proliferation to cell differentiation and expansion [6]. In plant roots, cells at the tip divide actively by normal mitosis, while endocycling cells become frequent further from the tip, in a zone associated with differentiation and increases in cell size [7, 8].

We developed a system to analyze DNA replication in *Zea mays* (maize) roots [8, 9], with similar approaches being applied in our work with *Arabidopsis* cell suspensions [10]. In this system, newly replicated DNA is labeled *in vivo* with the thymidine analog, 5-ethynyl-2’-deoxyuridine (EdU), and labeled nuclei are separated by flow cytometry into populations representing different stages of S phase. Cytological analysis showed that spatiotemporal features of maize DNA replication are significantly different from those of animal cells [11]. We then characterized the replication timing (RT) program in mitotic cells of the apical 1-mm root segment [12], using a modified replication timing by sequencing protocol (Repli-seq) [13, 14]. In mitotic cells, we found evidence for a gradient of early replicating, open chromatin that transitions gradually into less open and less transcriptionally active chromatin replicating in mid S phase. We also confirmed previous cytological observations showing that heavily compacted classical heterochromatin, including knobs and pericentromeres, replicate primarily in late S phase [11, 15]. While these relationships between RT and chromatin packaging are generally similar to those found in other systems, we did not find evidence for megabase-scale replication domains that have been characterized in mammalian cells (reviewed in [16] and references therein).

Although replication in the first 1-mm of the root is mostly mitotic, with DNA contents of labeled nuclei ranging from 2C to 4C, flow cytometry profiles of nuclei derived from root tissue between 1 and 3 mm from the tip also included a substantial population of labeled nuclei from endocycling cells, with DNA contents between 4C and 8C. Cytological analysis showed that the spatiotemporal patterns of replication in endocycling nuclei are very similar to those in mitotic nuclei [11]. However, it remained to be determined whether the entire genome is uniformly replicated during the endocycle, and whether the temporal program is altered when replication occurs without an intervening mitosis.

Both under-replication and over-replication (amplification) have been observed in multiple animal systems, notably including *Drosophila* (reviewed in [17]). In addition to the well-known amplification of chorion genes and under-replication of heterochromatin, under-replication also occurs in a number of euchromatic regions, with a degree of tissue specificity suggesting a possible role in differentiation [18–20].

Even though endopolyploidy is common in plants, there are very few reports of over- or under-replication of specific sequences. Some orchids exhibit a phenomenon in which only a fraction of the genome is endoreplicated [21, 22], but in most cases, endopolyploid cells have DNA contents that are multiples of the 2C value. Both highly repetitive heterochromatic regions and highly expressed genes are extensively endoreduplicated in maize endosperm nuclei, as would be expected for uniform replication of the entire genome [23]. More definitively, whole genome sequencing in *Arabidopsis* showed that leaf nuclear DNA is evenly endoreduplicated in wild-type plants, although the same series of experiments clearly demonstrated selective over-replication in *atxr5* and *atxr6* mutants [24].

In addition, there is as yet no information as to whether changes in RT programs are associated with endoreduplication or differentiation in plant systems. That such changes might occur in association with differentiation is supported by reports of extensive changes in RT between animal cell cultures representing different embryonic or differentiated cell types (e.g. [13, 25–27]).

To address these questions in the maize root tip system, we carried out a detailed comparison of RT dynamics in mitotic and endocycling cells. To isolate endocycling nuclei, we focused on a root segment 1–3 mm from the apex where there is a higher proportion of endocycling cells and used flow cytometry to separate nuclei of higher ploidy. We found very little evidence for changes in copy number that would be associated with over- or under-replication, and the RT profiles for the vast majority of the genome are very similar. However, we found significant changes in timing for a number of loci that together correspond to 2% of the genome. Most notably, we found major changes in the RT of centromeres, which replicate mainly during mid S phase in mitotic cells, but primarily in late S phase of the endocycle.

## RESULTS

### Separating endocycling from mitotic nuclei

As reported previously and described in Methods, we used a 20-min pulse of the thymidine analog, EdU, to label newly replicated DNA in intact maize roots. This was followed by formaldehyde fixation and isolation of nuclei from defined segments of root tips (Fig 1A). Incorporated EdU was conjugated with Alexa Fluor 488 (AF-488) by “click” chemistry [28]. The nuclei were then stained with DAPI and fractionated by two-color fluorescence activated flow sorting to generate populations at different stages of the mitotic cell cycle or the endocycle [8, 9]. Fig 1B and 1C show flow cytometry profiles obtained for root segments 0–1 mm and 1–3 mm from the tip, respectively. Fluorescent signals from nuclei that incorporated EdU during S phase of a normal mitosis form an “arc” between 2C and 4C DNA contents, while nuclei labeled during the endocycle S phase form a similar arc between 4C and 8C. As seen in Fig 1C, the endocycle arc is more prominent in nuclei preparations from 1–3 mm root segments. To analyze endocycle RT, which we will describe in detail below, we separated labeled nuclei representing early, mid, and late S-phase fractions using the sorting gates shown in Fig 1C, adjusting the endocycle early gate to avoid contamination with mitotic nuclei in late S phase. Reanalysis of the sorted nuclei confirmed that there was good separation between the nuclei populations from the adjusted early sorting gate and the mid sorting gate (S1 Fig). The flexibility of the EdU labeling and flow sorting system also allowed us to collect unlabeled nuclei, representing non S-phase cells with 2C, 4C and 8C DNA contents. These nuclei were used to characterize selected histone marks following mitotic or endocycle replication and to investigate the copy number of individual loci across the genome.

**Fig 1.**
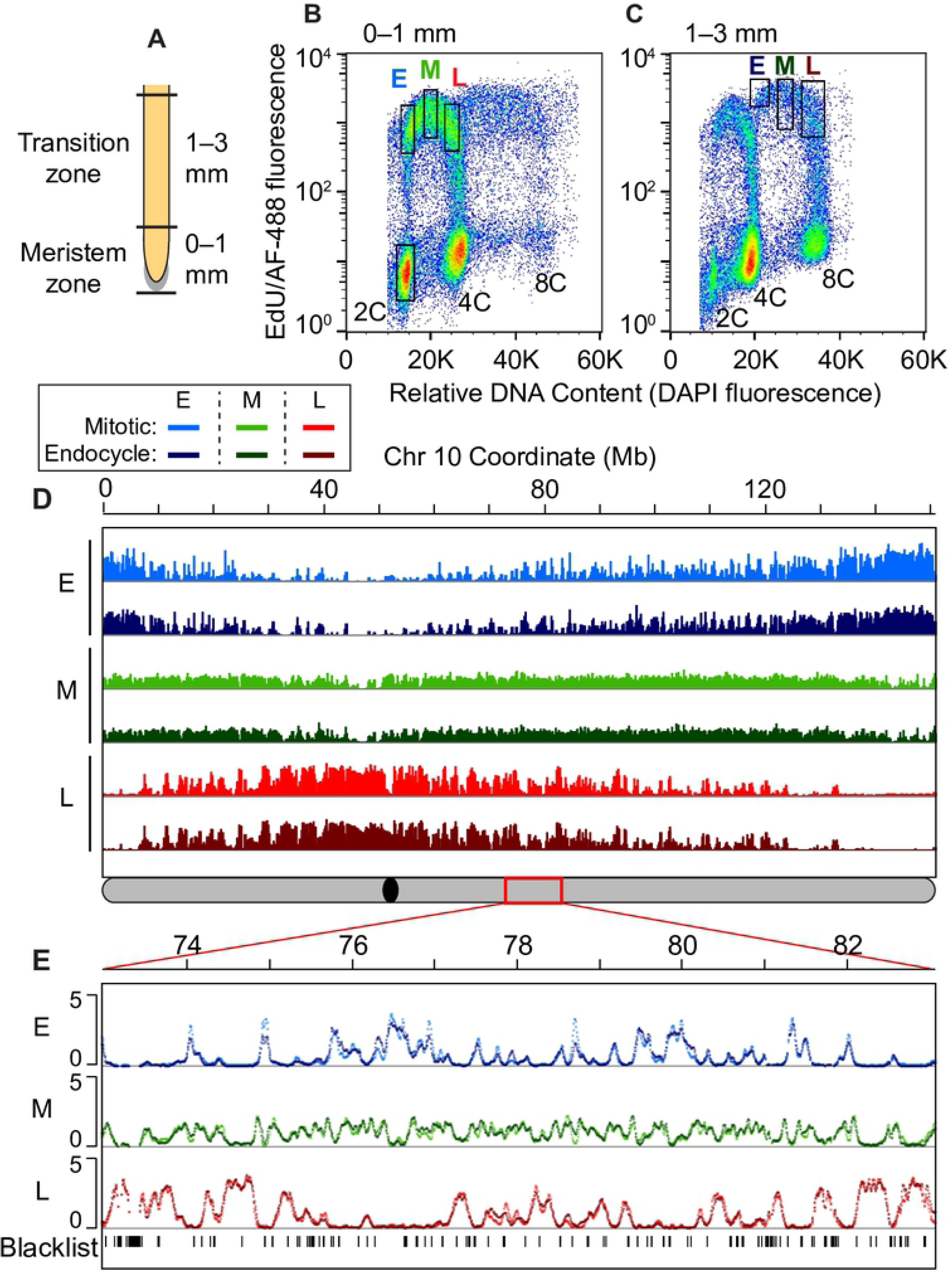
Global comparison of mitotic cycle and endocycle replication timing programs. (**A**) Schematic of a maize root showing the meristem zone (0–1 mm region) and transition zone (1–3 mm region) used for replication timing experiments. (**B and C**) Flow cytograms of nuclei isolated from the 0–1 mm root segments (**B**) and 1–3 mm root segments (**C**). Dots are pseudo-colored by density and black rectangles represent the sorting gates used to collect the pre-replicative 2C reference sample and early (E), mid (M) and late (L) S-phase fractions from either the mitotic cycle or endocycle. (**D**) Global scale view of replication timing (RT) for chromosome 10, comparing mitotic and endocycling profiles in early, mid and late S phase. Uniquely mapping reads were aggregated in 3-kb windows, normalized for sequencing depth, divided by the normalized 2C reference read counts, and Haar wavelet smoothed (see Methods). The global RT profiles for mitotic and endocycling cells are very similar to each other for all ten chromosomes. The schematic of chromosome 10 at the bottom shows the location of the centromere (black oval) and the 10 Mb region that is expanded in panel **E** (red rectangle). (**E**) Expanded view of a 10 Mb region on chromosome 10 with overlaid mitotic and endocycle RT profiles. Unmappable or multi-mapping regions (“blacklist”) are indicated as tick marks in the bottom track. This example illustrates the similarity between the mitotic and endocycle RT profiles that is observed throughout most of the genome. Scale for all panels: 0–5 normalized signal ratio.

### Evidence for complete genome replication during the endocycle

Given the well documented examples of over- and under-replication during the endocycle in animal systems, we investigated whether there are local copy number differences in the maize genome after endocycle replication. To do this, we used the non S-phase 2C, 4C, and 8C nuclei populations described above, and carried out whole genome paired-end sequencing. To gain a better representation of the copy number of repeat regions in the genome, reads that could not be uniquely mapped to a single location were included, but we retained only the primary alignment location for each read pair. These data were examined for regions in which normalized read frequencies in 5-kb windows differed between 8C and 4C or 4C and 2C nuclei, using procedures described by Yarosh et al. ([29]; S1 Text). We found about 5% of the 5-kb windows had ratio values that fell outside of two standard deviations of the mean ratio for 8C and 4C or 4C and 2C (1.0 ± 0.2 S. D. for both; S2A and B Fig). However, these windows all either occurred as singleton 5-kb windows scattered around the genome (S2C Fig) or coincided with regions that had very low read mapping in the 2C sample, indicating they are likely the spurious result of making a ratio between windows with very few reads in both samples. As such, there is very little evidence of meaningful over- or under-replication of genomic regions in nuclei with different ploidy levels.

To further investigate whether there is complete replication of high-copy repeats that are not well represented in the genome assembly, we used BLAST software to query all reads, not just those that can be mapped to the genome, to determine the percentage of reads corresponding to each of several consensus sequences for high-copy repeats (S1 Text). Analyzed sequences included the knob repeats *knob180* and *TR-1* [30, 31], 5S and 45S rDNA repeats [32], and centromere-associated *CentC* satellite repeats [33]. We also queried consensus sequences for centromere retrotransposons of maize (*CRM*) families 1–4 [34–37]. In all cases, we found the percentages to be similar in the 2C, 4C and 8C samples (S2D and E Fig), further suggesting that there is little or no over- or under-replication.

### Replication timing analysis

As described above, we sorted endocycling nuclei from the S-phase populations in Fig 1C, and extracted and sheared the DNA in each fraction. EdU-containing DNA fragments were immunoprecipitated with an antibody to AF-488, resulting in sequence populations representing DNA replicating during early, middle, or late S phase of the endocycle. We also prepared DNA from the unlabeled 2C nuclei pool to provide a reference dataset representing pre-replicative nuclei. DNA from three biological replicates of each sample was sequenced to generate paired-end reads.

To compare the RT programs in endocycling and mitotic nuclei, we mapped our previous Repli-seq data for mitotic nuclei [12] and our new data for endocycling nuclei to the new maize B73 RefGen_v4 genome, which includes improved assemblies of centromeres and more complete annotations of transposable elements (TEs) [38, 39]. Uniquely mapped read depth varied between ∼3 and 11× genome coverage per S-phase sample, so all samples were randomly downsampled to ∼3× coverage to ensure comparable results (see Methods and S1 Spreadsheet).

We used the *Repliscan* analysis pipeline [14] to generate profiles of replication activity in early, mid and late fractions of each S phase. These profiles were generated by aggregating the Repli-seq read densities for each S-phase sample in 3-kb static windows, scaling the reads to 1× genome coverage, and then dividing by the scaled read counts from the unlabeled 2C reference data and smoothing by Haar wavelet transform (see Methods and [14]). Normalizing with the 2C reference corrected for differences in sequencing efficiencies and collapsed repeats that caused “spikes” in the data (illustrated for late replication in the endocycle in S3 Fig), producing an estimate of replication intensity or “signal” in each 3-kb window. We also excluded 3-kb windows with extremely low read coverage in the 2C reference sample (see Methods) from all analyses (“blacklist” windows, indicated by black tick marks in Fig 1E).

Fig 1D shows that the global RT patterns are remarkably similar in endocycling and mitotic nuclei, and overlays of the corresponding profiles show mostly minor differences (Fig 1E). Pearson’s correlation coefficient values between corresponding S-phase fractions from the mitotic and endocycle data are very high (r values of 0.91, 0.89 and 0.96 for early, mid and late, respectively). These values are similar to those found between individual biological replicates within each sample (S4 Fig).

### Identifying regions of altered timing

Despite the global similarity of the RT programs of mitotic and endocycling cells, there are regions scattered around the maize genome that show a shift in RT. To identify timing differences, we first calculated the difference in normalized replication signal between the mitotic and endocycle data at each genomic location for the early, mid and late profiles separately (S1 Table; S5 Fig). We then constrained our analysis by focusing only on regions where there was an equal and opposite timing difference in at least one other S-phase fraction (for example, regions in which a decrease in early replication signal in endocycling cells was associated with a corresponding increase in mid and/or late S-phase signal at the same location). We allowed a gap distance of 6 kb when searching for regions with timing differences to account for small blacklist regions that break up larger regions of change. We found that 11% of the genome showed a difference in timing of at least 10% of the total difference range for a given profile (difference in replication signal ≥ 0.4; S1 Table), with an opposite timing difference at the same threshold criterion at the identical location in another S phase profile. Many of these regions are small, with the lower 50% of regions ranging in size from 3 kb to the median size of 33 kb (S2 Table), and it is not clear if such small alterations are biologically relevant.

To identify more robust differences, designated <u>R</u>egions of <u>A</u>ltered <u>T</u>iming (RATs), we identified regions in which the difference in replication signal was ≥ 25% of the total difference range for a given profile (difference in replication signal ≥ 1.0; S1 Table), and which also met the criterion of having an opposite difference in at least one other profile. To highlight larger and contiguous regions of change, we included ≥ 10% regions that were adjacent to the original ≥ 25% regions. However, RATs had to have at least one core region where the timing change was at least 25% (S2 Table) to be included in our analysis. Representative ≥ 25% and ≥ 10% regions are indicated by different shades of red and blue bars in Fig 2 (additional examples are in S6 Fig). Finally, we examined the profiles for the RATs in individual biological replicates to verify there was good agreement between the replicates (Figs 2B and S6). By selecting only the most robust RATs we excluded other regions where timing changes are less dramatic – for example those indicated by dashed boxes in Fig 2. In such regions, the timing difference did not meet our criteria of a ≥ 25% difference in signal (box 2 in Fig 2A) and/or there is not an equal and opposite (“compensated”) timing difference (box 3 in Fig 2A).

**Fig 2.**
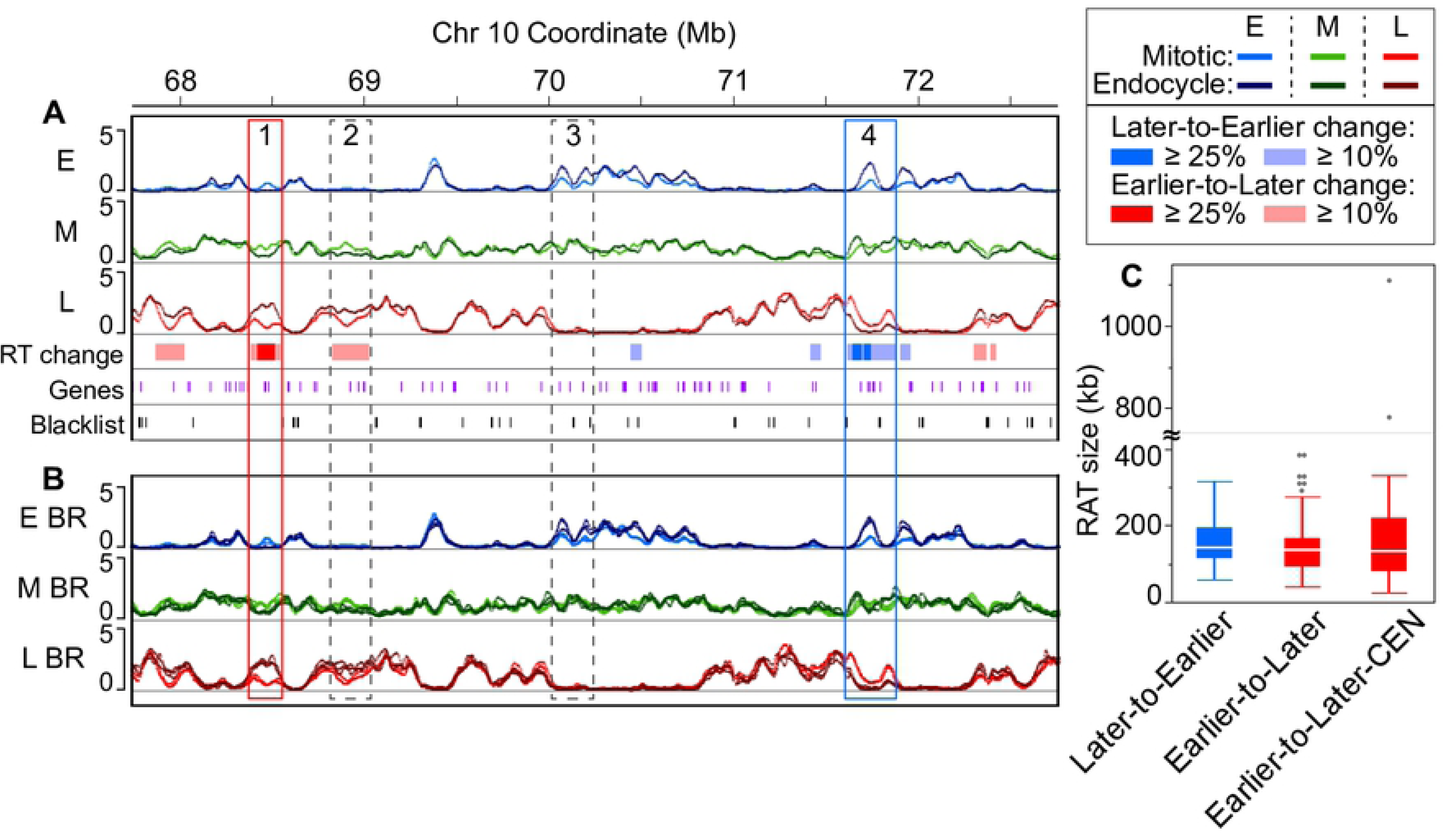
Identifying regions of altered timing. (**A**) An example region (5 Mb) on chromosome 10 containing two robust Regions of Altered Timing (RATs), indicated by boxes outlined with solid lines. The RAT in box 1 (red) shifts from Earlier-to-Later, and the RAT in box 4 (blue) shifts from Later-to-Earlier. Dashed boxes denote regions with some level of RT difference in which the magnitude of the difference did not meet our ≥ 25% criterion (box 2), or in which the change in one S-phase fraction was not compensated by an opposite change in at least one other S-phase fraction (box 3). Annotated genes (purple) and unmappable or multi-mapping regions (“blacklist”, black) are indicated as tick marks in the bottom tracks. (**B**) The same chromosome region as in (**A**) with the individual biological replicate profiles overlaid to demonstrate that RATs are not caused by local regions of technical variation between replicates. Scale for panels **A** and **B**: 0–5 normalized signal ratio. (**C**) Boxplots representing the distribution of RAT sizes in the three categories: Later-to-Earlier, Earlier-to-Later, and a subset of Earlier-to-Later RATs found in functional centromeres (CEN) [38]. Boxplot whiskers represent 1.5 x interquartile range (IQR). The axis is broken to show two values that are much higher than the others and correspond to large RATs in CEN 9 and CEN 10. However, it is important to note that the sizes of CEN RATs are underestimated, because centromeres contain variable numbers and sizes of blacklist regions, which break up what would probably be long continuous RATs (see Fig 3).

Robust RATs fall into two categories, those where the strongest replication signal occurs later in the mitotic cycle than it does in the endocycle (“Later-to-Earlier” shift), and those in which the strongest signal occurs earlier in the mitotic cycle than in the endocycle (“Earlier-to-Later” shift). In addition, we separately characterized a subset of the Earlier-to-Later RATs that are located in functional centromeres (“Earlier-to-Later-CEN”) using centromere (CEN) coordinates from [38]. Our stringent criteria identified RATs comprising only about 2% of the maize genome (Table 1), with the vast majority (1.7% of the genome) in the Earlier-to-Later category. Non-CEN Later-to-Earlier and Earlier-to-Later RATs have similar size distributions, with median sizes of 141 and 135 kb, respectively (Fig 2C and Table 1). All of the CEN RATs fall into the Earlier-to-Later category and have a median size of 132 kb, similar to the non-CEN RATs. It is important to note, however, that the sizes of CEN RATs are likely underestimated because of numerous blacklist regions within the centromeres that break what are likely continuous RATs into several smaller parts in our analysis. Even though maize centromeres are remarkably well sequenced [38], they still contain some gaps and regions where reads cannot be uniquely mapped in the current B73 RefGen_v4 genome assembly, as indicated by the black tick marks in the bottom tracks of Fig 3A–3D.

**Fig 3.**
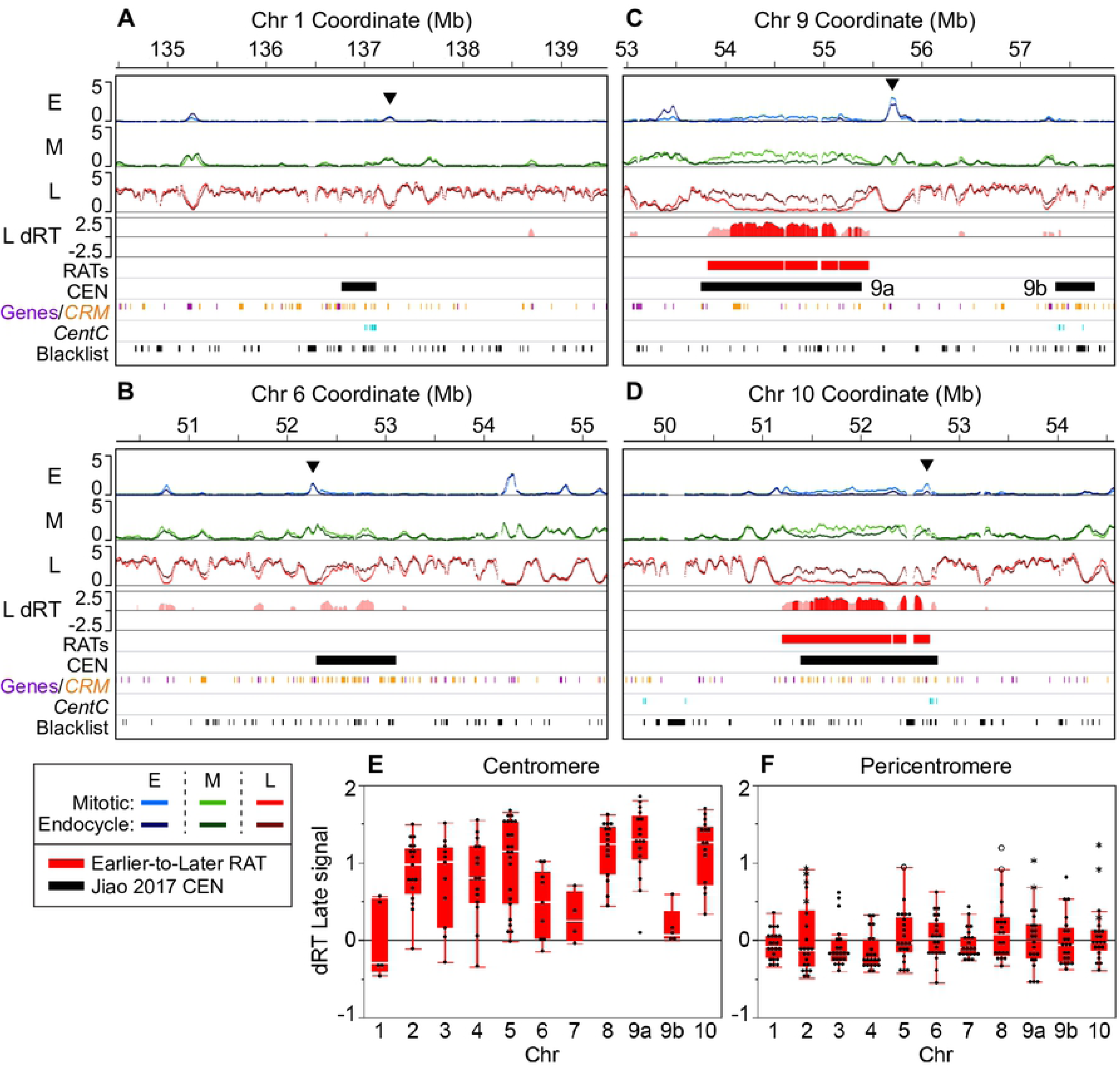
Large RATs correspond to functional centromeres. Our analysis found large RATs, sometimes broken by blacklist regions (black tick marks at the bottom of each panel) at each of the seven “complex” maize centromeres. The remaining three “simple” centromeres (on chromosomes 1, 6, and 7) showed various levels of timing differences that did not meet the criteria for calling RATs in our initial analysis. (**A–D**) Each 5-Mb region shown contains early (E), mid (M) and late (L) RT profiles with mitotic and endocycle data overlaid (scale: 0–5 normalized signal ratio). The difference in late replication signal profiles (endocycle minus mitotic; labeled “L dRT”) for windows where the difference was compensated by an equal and opposite difference in the early and/or mid profiles is also shown. Late differences compensated at the ≥ 10% threshold (light red), and those compensated at the ≥ 25% threshold (dark red) are shown, but only regions that contained at least one ≥ 25% shift were classified as robust RATs in our initial analysis. Two examples of simple centromeres, CEN 1 (**A**) and CEN 6 (**B**), and two examples of complex centromeres, CEN 9 (**C**) and CEN 10 (**D**) are presented. The black arrowheads in panels **A–D** denote example regions with a peak of early replication signal within or adjacent to the centromere (for other examples, see S12 Fig). Colored boxes below the RT profiles denote Earlier-to-Later RATs (red) and the functional centromere (black; [38]). Chromosome 9 contains two called CEN regions labeled 9a and 9b. The colored tick marks (see legend for colors) correspond to elements of centromeric retrotransposons of maize (*CRM*) families 1–4 [39], gene annotations in RefGen_v4 [38] and the locations of mappable *CentC* satellite repeats [40]. Blacklist regions are indicated by black tick marks in the lowest track. (**E and F**) Timing differences (endocycle - mitotic) between late profiles for each centromere (**E**) and corresponding pericentromere (**F**; ± 1 Mb) were calculated in 100-kb static windows. In panel **F**, asterisks indicate difference values from windows where an Earlier-to-Later-CEN RAT extends past the called CEN boundary [38] into the pericentromere; open circles indicate windows that contain a non-CEN Earlier-to-Later RAT that met our compensation criteria. Timing differences between early and mid profiles are shown in S13 Fig.

**Table 1.** RAT summary table.

| RAT category | Count | Median size (kb) | Coverage (kb) | % of genome | RATs with gene (%) | RATs with expressed gene (%) |
| --- | --- | --- | --- | --- | --- | --- |
| Later-to-Earlier | 41 | 141 | 6,291 | 0.3 | 92.7 | 82.9 |
| Earlier-to-Later | 192 | 135 | 26,907 | 1.3 | 96.4 * | 91.1 * |
| Earlier-to-Later-CEN | 41 | 132 | 7,668 | 0.4 | 43.9 | 22.0 |

A summary of the region count, median size, total genome coverage, and percentage of the entire genome represented in each RAT category. The number of RATs that overlap genes or expressed genes is also presented. Asterisks denote one RAT category in which the indicated percent overlap was greater than expected by chance (permutation *P* value ≤ 0.001), estimated by permutation analysis (see Methods and S7 Fig.).

### Non-centromeric RATs

We analyzed the non-CEN RATs for the content of genes and TEs, as well as the presence of histone modifications and functional annotations related to the genes within RATs. To assess whether the percentage of RATs containing genes differed from random expectation, we randomly shuffled coordinates corresponding to the non-CEN Later-to-Earlier and Earlier-to-Later RATs around the genome 1000 times and calculated the percentage of regions that overlap genes in each set. We found that 93% and 96% of Later-to-Earlier and Earlier-to-Later RATs, respectively, contain at least one annotated gene and usually contain a small cluster of genes (Tables 1 and S3). Using root-tip RNA-seq data that are not specific to mitotic or endocycle cells, we found that although only 50% of the 682 genes found in non-CEN RATs are expressed at a meaningful level (FPKM ≥ 1; S3 Table), 83% and 91% of Later-to-Earlier and Earlier-to-Later RATs, respectively, contain at least one expressed gene (Table 1). The observed percent overlap of Earlier-to-Later RATs with genes and expressed genes are both significantly greater than expected by random chance (permutation *P* value ≤ 0.001; S7B and D Fig). Differences from random expectation were less obvious for Later-to-Earlier RATs, although the percent overlap of expressed genes is on the edge of significance (permutation *P* value = 0.035; S7C Fig).

We were unable to directly compare expression of genes in RATs in mitotic and endocycling cells because we could not obtain RNA of sufficient quality to sequence from fixed, sorted nuclei. Instead, we assessed a selection of gene-associated histone post-translational modifications in sorted non S-phase 2C, 4C and 8C nuclei. In our previous work in maize root mitotic cells, we showed that trimethylation of H3 lysine 4 (H3K4me3) and acetylation of H3 lysine 56 (H3K56ac) modifications tend to colocalize on active genes and are associated with earlier replicating regions, while trimethylation of H3 lysine 27 (H3K27me3) tends to be on repressed genes regardless of their RT [12]. For each ploidy level, we quantified the percentage of genes within RATs that have each mark, as well as the fold enrichment relative to input for called peaks within genes. There are very few differences between ploidy levels in the number of genes bearing these marks (S8D Fig), but there are some minor shifts in the peak enrichment in 8C nuclei compared to 2C (S8A–C Fig). The clearest shift is a decrease in H3K4me3 enrichment found on expressed genes in Earlier-to-Later RATs (S8B Fig), which suggests these genes may have decreased expression in endocycling cells.

We also performed a gene ontology (GO) analysis for the genes found in non-CEN RATs to ask if there are functional annotations enriched in genes that shift replication timing. For this analysis, we focused on the genes that we identified as expressed in the root tip (S2 Spreadsheet). We found 44 significantly enriched GO terms for genes within Earlier-to-Later RATs, including biological process and molecular function terms related to gene expression, DNA/RNA metabolism, and the cell cycle (S9 Fig). A wide variety of significant cellular component GO terms were also found, which may relate to various differentiation processes occurring in endocycling cells. There are no significant GO terms for genes within Later-to-Earlier RATs, though the presence of only 52 expressed genes in this RAT category made it difficult to fully assess significance. Taken together, these analyses of transcription-related histone modifications and functional annotations suggest a role for gene expression changes in the Earlier-to-Later RATs. Given that these regions are shifting to a later RT in the endocycle, a decrease in gene expression would be expected [12]. Clearly, however, more work will be needed to confirm this hypothesis.

The general organization of the maize genome is genes clustered in “islands” interspersed with blocks of transposable elements [41–43]. We used a similar permutation strategy as for the genes to estimate the significance of any differences in percent coverage of each TE superfamily in non-CEN RATs as compared to random expectation, estimated from 1000 randomly shuffled sets. The TE annotations were from the recent RefGen_v4 TEv2 disjoined annotation, where every bp is assigned to a single TE [39]. We found the coverage of the RLG/Gypsy superfamily in Earlier-to-Later RATs is significantly less than random expectation (permutation *P* value ≤ 0.001; S4 Table). There are other, less significant, positive and negative associations with TE superfamilies in non-CEN RATs, including RLC/Copia, DTT/Tc1-Mariner, DTM/Mutator and DHH/Helitron (S4 Table). We also found that the percent AT content in RATs is similar to that of the genome as a whole, with median values of 55% and 56% for Later-to-Earlier and Earlier-to-Later RATs, respectively, and a median value of 55% for the whole genome (S10 Fig).

### Centromeric RATs

Functional centromeres are defined by their content of nucleosomes containing the centromere-specific histone variant known as CENH3 in plants and CENP-A in animals. CENH3/CENP-A makes up only a small percentage of the total H3 population in centromeres, but plays an important role in recruiting kinetochore proteins [44–46]. Maize is unusual among higher eukaryotes in that a majority of centromeric reads can be uniquely mapped [47]. In our replication timing data, for example, we found that on average 45% of all reads that map to centromeres could be uniquely mapped to a single location (S11 Fig). Only these uniquely mapping reads were used for further analysis. In addition, most of the maize centromere assemblies are relatively intact, and functional centromeres have been located by mapping ChIP-seq reads for CENH3 [38]. When combined with our replication timing data, these features of the maize system create a unique opportunity to assess RT programs for centromeres.

Our analysis found large, robust RATs across seven of the ten centromeres (Figs 3C, 3D and S12), with replication occurring mainly in mid S in mitotic cells, but changing to primarily late S in endocycling cells. It is also noteworthy that though replication occurs mainly in mid S in mitotic cells, there are some distinct peaks of early replication inside or directly adjacent to the called centromere (indicated by black arrowheads in Fig 3 and S12) in all but one of the maize centromeres. These early peaks remain in the endocycle, though in some cases there is a reduction in early signal with a concomitant increase in mid signal at the same location. The seven centromeres that contain robust RATs (CEN 2, 3, 4, 5, 8, 9 and 10) were previously classified as “complex” because they contain a mixture of retrotransposons with some centromere satellite repeat arrays (*CentC*; [40, 47]). In the RefGen_v4 genome assembly, CEN 9 has two called CENH3-binding regions [38], which we refer to as CEN 9a and 9b (Fig 3C; black bars). Interestingly, we only found a robust RAT in the larger CEN 9a, with the smaller CEN 9b showing almost no timing shift.

The remaining three centromeres (CEN 1, 6, and 7) were previously characterized as “simple” because they mainly contain large arrays of the *CentC* repeat [40, 47]. In our analysis, the simple centromeres showed, at most, small timing shifts that did not meet our criteria for a robust RAT (Figs 3A, 3B and S12). However, *CentC* repeats are not well represented in the reference genome assembly, so our ability to analyze replication of the complete simple centromeres is limited. Portions of CEN 7 that are present in the assembly replicate mainly in mid S phase in both mitotic and endocycling cells (S12 Fig), while sequences in the assemblies for CEN 1 and CEN 6 are mostly late replicating in both types of cells, with some minor timing changes across small regions (Fig 3A and 3B).

The robust RATs on the seven complex centromeres correspond quite closely to the boundaries of the functional centromeres defined from CENH3 ChIP-seq data [38]. The cumulative coverage of RATs in each complex centromere ranges from 405–1518 kb (S5 Table). However, because each centromere includes blacklist regions that vary in size and number, automated analysis did not identify the true sizes of the RATs. To avoid this problem, we have chosen to focus the following analyses on the entire functional centromere instead of on computationally identified RATs.

For the entire CENH3-binding region of each chromosome (excluding blacklist regions), we calculated the difference in early, mid and late replication signal (endocycle minus mitotic) from RT profiles by averaging across 100-kb static windows. For comparison, we also calculated the replication signal differences in pericentromeres, which were arbitrarily defined as the ± 1 Mb flanking the CENH3 region. We inspected all RT differences in the centromeres and pericentromeres by not requiring that the RT differences be compensated by an opposite shift in the other S-phase fractions. Early and mid replication signals across the complex centromeres decrease and late replication signals increase in endocycling cells, reflecting a large shift toward late replication. The RT difference values for the late profile in centromeres and pericentromeres are shown in Fig 3E and 3F, respectively, while the difference values for early and mid profiles are shown in S13 Fig. Interestingly, the timing difference tapers off towards the edges of the functional centromere (see profiles in Figs 3C, 3D and S12), and there is striking congruity in the replication signals for mitotic and endocycling cells in the immediately adjacent pericentromere regions (Fig 3A–D). The few timing shifts in pericentromeric regions are smaller in size and much less dramatic than those in the centromere proper (Fig 3F). Moreover, very few (8%) of pericentromeric windows with timing shifts are compensated by an equal and opposite shift in the other S-phase profiles (S6 Table), suggesting many of these uncompensated differences may result from technical variation rather than from meaningful biological differences. In contrast, nearly all (85%) of the centromeric windows have compensated RT shifts.

### Genomic elements and features in centromeres

Maize centromeres contain varying amounts of tandemly arrayed *CentC* repeats (single repeats of 156 bp in length; [33]) as well as several *CRM* retrotransposon families interspersed with elements from a few other retrotransposon families [36, 43, 48, 49]. *CentC* repeats and *CRM* elements are also present in the adjacent pericentromeres where there is no CENH3 binding [43, 48]. In RefGen_v4, there are also fifty annotated genes within centromeres. We asked if all of these sequence elements in centromeres behave similarly in the mitotic to endocycle transition, or if certain elements show larger timing shifts than others. We also asked if all three types of sequence elements show similar RT changes in centromeres versus pericentromeres. Given that the RT signal values were aggregated in 3-kb windows, we only included elements that covered at least half a window (1.5 kb) in our analysis. Fig 4 summarizes data on these questions for the complex centromeres, while data for the simple centromeres are shown in S14 Fig. Similar results were found when all elements were included (S14 Fig).

**Fig 4.**
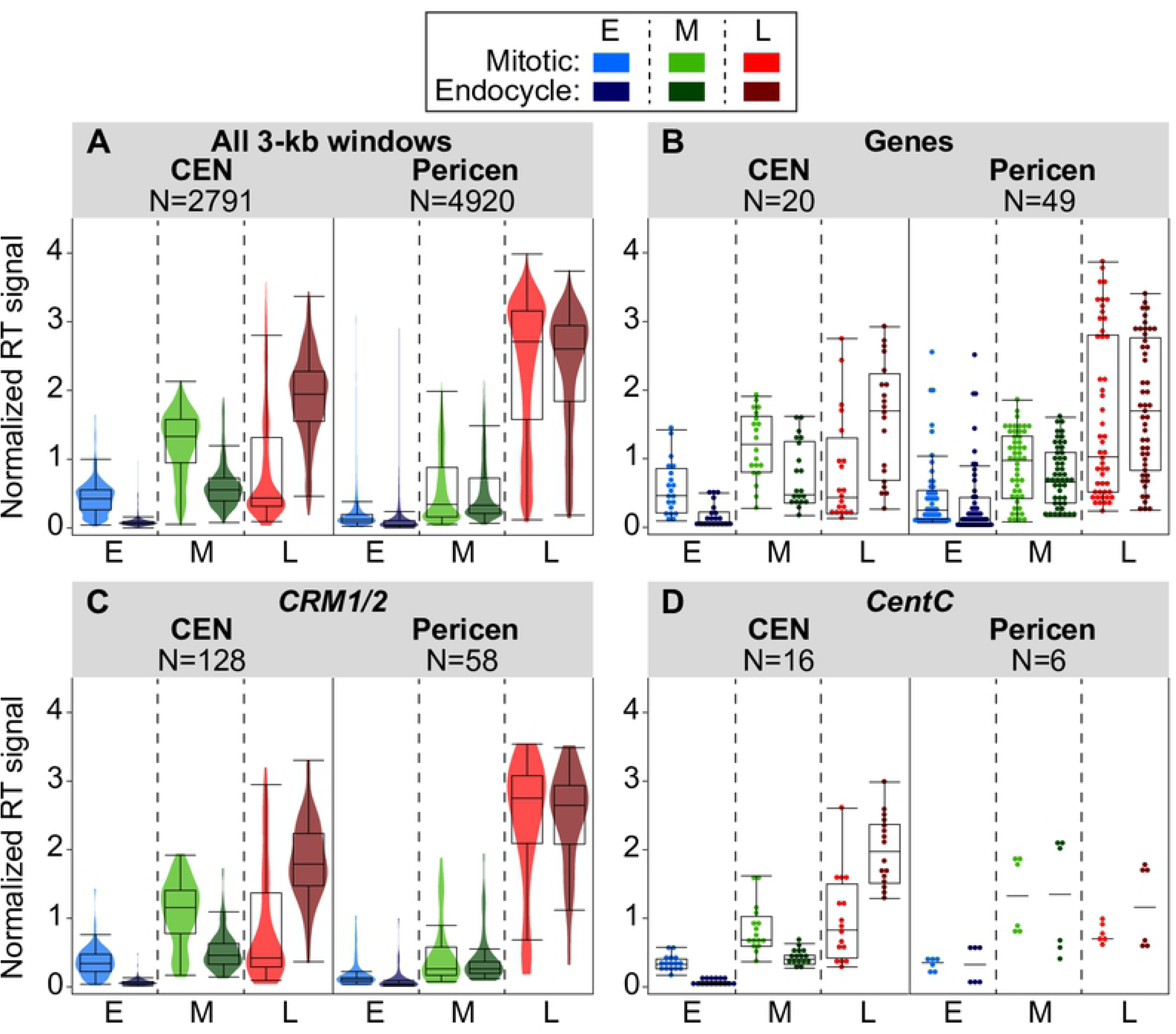
Comparing replication times for genomic features in complex centromeres and corresponding pericentromeres. **(A–D)** Boxplots comparing replication signals during mitotic and endocycle S phases for centromeres, pericentromeres (± 1 Mb), and genomic features within them. The panels show the distributions of replication signals in early (E), mid (M), and late (L) S for all 3-kb windows (**A**), annotated genes (**B**), mapped *CentC* repeats (**C**), and *CRM1/2* elements (**D**) in centromeres and pericentromeres. For panels **A** and **C**, colored violin plots are overlaid, while for panels **B** and **D**, individual data points are shown. Only elements that covered at least 50% of a 3-kb window were included in each analysis, though results were similar when all elements were included (S14 Fig). The number of windows or elements included in each analysis is indicated above each graph. Boxplots for all elements in simple centromeres, as well as for the individual *CRM1* and *CRM2* families are in S14 Fig.

The results for the two dominant *CRM* families, *CRM1* and *CRM2*, are similar (S14 Fig), so these families were grouped together in Fig 4C. When present in centromeres, all three major classes of elements – genes, *CRM1/2*, and *CentC* repeats – clearly replicate later during the endocycle than in the mitotic cycle (Fig 4). In contrast, genes and *CRM* elements in the pericentromere show little or no timing shifts. A full analysis of the replication times of *CentC* repeats in pericentromeres is hampered by the limited representation of this repeat class in the genome assembly (Fig 4D and S14E).

### Chromatin features in centromeres

We also examined activating (H3K56ac and H3K4me3) and repressive (H3K27me3) histone post-translational modifications to look for epigenetic changes in centromeres after endocycle replication. It was previously reported that some H3K4me3 and H3K27me3 peaks of enrichment occur in the centromere, mainly associated with genes [50]. We asked whether genes that have these modifications continue to have them after mitotic and endocycle replication, and found very few changes in the number of genes with these modifications at each ploidy level (S15 Fig). There was also very little change in the fold enrichment of these histone marks in centromere genes when comparing 2C, 4C and 8C nuclei.

We also investigated the levels of dimethylation of histone H3 lysine 9 (H3K9me2) enrichment in each centromere. Previous work indicated there is a depletion of H3K9me2 in centromeres relative to adjacent pericentromeres [51, 52], which we observed as well (S16 Fig). Traditional peak calling tools are not effective for H3K9me2 because of its even distribution across the maize genome. Instead, we estimated the fold enrichment by calculating the percent of total H3K9me2 ChIP reads in a given centromere region (using coordinates from [38]) and dividing by the percent of total input reads corresponding to that centromere in three biological replicates). We found a similar H3K9me2 average fold enrichment for all centromeres and for 2C, 4C and 8C nuclei, although values for 4C and 8C nuclei were consistently slightly higher than those for 2C nuclei (S16A Fig). CENH3 nucleosomes lack the lysine 9 residue found in canonical histone H3 [53], so H3K9me2 enrichment must occur in the interspersed H3 nucleosomes.

### Centromeric histone H3 in mitotic and endocycling centromeres

Unlike the canonical histone H3, CENH3 is not replaced in a replication dependent manner in higher eukaryotes, resulting in a dilution of CENH3 relative to centromeric DNA during S phase [54, 55]. New CENH3 is incorporated into nucleosomes after the completion of S phase, but the timing of its integration into centromeric chromatin differs for plants, flies and humans (reviewed in [56]). In the plants tested thus far, deposition of CENH3 has been reported to occur between late G2 and metaphase [57–60].

Because mitosis does not occur in the endocycle and centromere function is presumably not required, we speculated that CENH3 might remain at low levels following DNA replication in endocycling cells. This hypothesis is supported by cytological studies of *Arabidopsis* endopolyploid nuclei showing the CENH3 signal does not increase in parallel with the total DNA content or the signal for 180-bp centromeric repeats [58, 59]. To test this hypothesis with maize centromeres, we used a maize anti-CENH3 antibody [48] for ChIP-seq analysis of CENH3 binding in sorted non S-phase 2C, 4C, and 8C populations of nuclei. It is important to note that the 4C nuclei come from a mixture of cells, some of which will return to the mitotic cycle and others that will continue on to the endocycle (at least 13% of nuclei in the 1–3 mm region). We asked whether the location or level of CENH3 enrichment changed after DNA replication in the mitotic cycle or the endocycle. For visualization of CENH3 localization, ChIP-seq read counts from three biological replicates for each ploidy level were aggregated in 3-kb windows and normalized to the level of a uniform 1× genome coverage, so that corresponding windows in the different ploidy level profiles were comparable. The normalized read count in each 3-kb window was then divided by the corresponding normalized read count for the corresponding ploidy input DNA to calculate a fold enrichment relative to DNA content value for CENH3 binding sequences in that window. The spatial distribution of CENH3 enrichment across the centromeres remained the same in 2C, 4C, and 8C cells. This is illustrated for CEN 9 and CEN 10 in Fig 5A and 5B, and data for the rest of the centromeres are shown in S17 Fig. There are also a few small spikes of CENH3 enrichment outside the called centromere (e.g. seen in Fig 5 and S17, but also occasionally further out on the arms). These spikes also remain in the same location between 2C, 4C and 8C cells, some of which could be related to misassembly of the reference genome. However, if real, these ectopic CENH3 peaks are less numerous and more persistent in G2 than those recently observed in HeLa cells [61].

**Fig 5.**
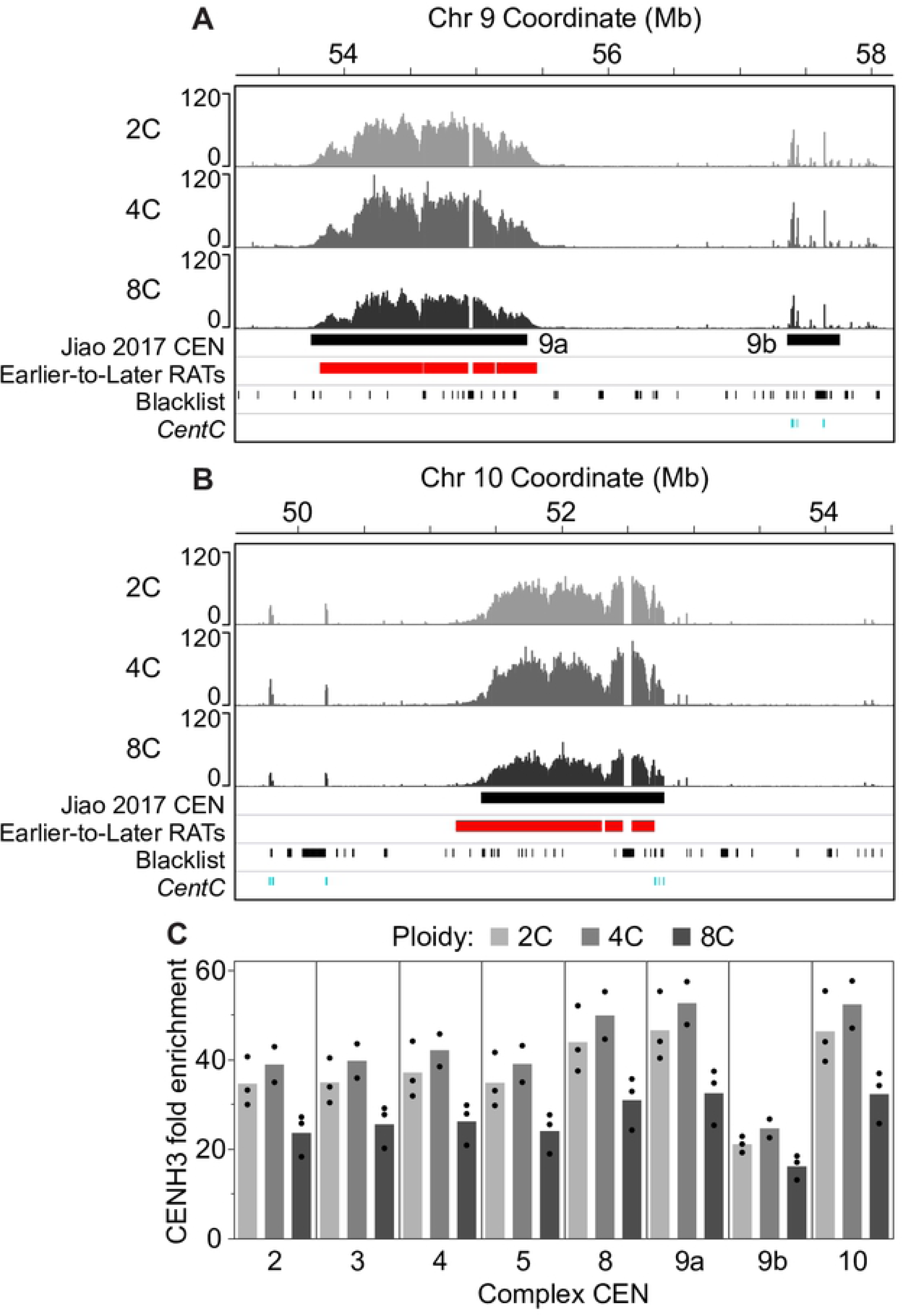
CENH3 localization and enrichment in mitotic and endocycling centromeres. We profiled CENH3 binding by ChIP-seq in flow sorted, non S-phase nuclei with 2C (before mitotic replication), 4C (after mitotic replication) and 8C (after endocycle replication) DNA contents. (**A and B**) CENH3 localization patterns for 2C, 4C and 8C nuclei in CEN 9a and 9b (**A**) and CEN 10 (**B**). Scale in both panels is 0–120 fold CENH3 enrichment relative to input. Colored boxes below the CENH3 profiles denote the previously identified functional centromere (black; [38]), and Earlier-to-Later-CEN RATs (red). Tick marks in the bottom two tracks indicate blacklist regions (black) and mapped *CentC* repeats (teal). (**C**) We used the ChIP-seq datasets from 2C, 4C and 8C nuclei to estimate the CENH3 average fold enrichment relative to DNA content for complex centromeres by calculating the percent of total CENH3 reads found in a given centromere (using coordinates from [38] and dividing by the percent of total input reads corresponding to that centromere. Black dots represent the individual values from biological replicates. Data for simple centromeres are shown in S17B Fig.

To compare total CENH3 content of entire centromeres at different ploidy levels, we calculated the percent of total CENH3 reads found in a given centromere and made a ratio to the percent of total reads from the corresponding input DNA in that centromere separately for each biological replicate, as described above for H3K9me2. The CENH3 average fold enrichment relative to total DNA content is similar for 2C and 4C nuclei in each of the complex centromeres (Fig 5C), with an average 4C/2C enrichment ratio of 1.1 (S7 Table). However, CENH3 enrichment decreases with the increase in ploidy from 4C to 8C (Fig 5C), with an average 8C to 4C enrichment ratio of only 0.7 (S7 Table). Average CENH3 enrichment values for simple centromeres were lower and slightly more variable, likely because of assembly issues. In both cases, however, the ratio of CENH3 enrichment in 8C cells to that in 4C cells is clearly higher than 0.5, which would be expected if there was no incorporation of new CENH3 after endocycle replication, but smaller than the 1.0 ratio expected if there was full replacement (S7 Table). It is worth noting that these data refer to post-replication 8C nuclei, which exited S phase prior to the time of analysis, and that post-replication 4C nuclei show no dilution of CENH3 relative to DNA content. Thus, our data are consistent with a model in which the CENH3 to DNA ratio is reduced as DNA replicates during the endocycle S phase, and only partially restored after completion of S phase.

## DISCUSSION

The maize root tip includes a naturally occurring developmental gradient, with cells in the meristem region (ca 0–1 mm) primarily undergoing mitotic cell cycles, while a subpopulation of cells in the transition zone (ca 1–3 mm) enters a developmentally programmed endocycle prior to further differentiation [8, 62]. Even though endocycling is very common in plants and plays essential roles in differentiation and the development of specialized tissues, cell size increases, and stress responses [2, 5, 63, 64], replication timing (RT) programs have not yet been characterized for alternative cell cycles, such as the endocycle.

We generated whole genome Repli-seq data for root cell nuclei undergoing DNA replication in either the mitotic cycle or the endocycle, making use of *in vivo* EdU labeling of intact root tips and two-color fluorescence activated nuclei sorting. By doing so, we avoided potential artefacts caused by cell synchronization [65] and chromosome aberrations often found in plant and animal cell cultures (e.g. [66–68]). We present replication activity profiles for early, mid and late replication separately, instead of collapsing the data into an early:late ratio as many studies do. The rationale for this approach is that, for roughly one third of the maize genome, we previously found heterogeneity in mitotic RT – e.g. regions of the genome in which root tip cells exhibit significant replication activity in both early and mid S, or both mid and late S [12]. An additional advantage to presenting the replication profiles separately is the ability to assess whether there are concomitant or “compensated” changes in a region at multiple stages of S phase. This compensation criterion helped us separate RT shifts that could be subject to technical error, such as alterations in flow sorting gates, from shifts that are more likely to represent meaningful changes in the population preference to replicate a replicon or cluster of replicons at a particular time in S phase.

The current study sought to investigate whether the mitotic RT program is maintained in the first round of the endocycle in maize root cells, despite the need to replicate twice as much DNA and the initiation of various root cell differentiation pathways. Extending our previous cytological observation that spatiotemporal patterns of replication are similar in mitotic and endocycling cells [11], we found that RT programs at the sequence level are strikingly similar as well. Pearson’s correlation coefficient values comparing data from the two types of cell cycles were similar to those for biological replicates within each type. The high level of reproducibility is particularly noteworthy in the case of the early replication profiles, given that the flow sorting gate for early replicating nuclei in the endocycle had to be adjusted to minimize contamination from late replicating mitotic nuclei (Fig 1C). This overall conservation of RT programs suggests that the process of re-establishing the RT program must be similar for the two types of cell cycles in maize roots. In animal systems, re-establishment of the RT program has been shown to occur in G1 of each cell cycle at a “timing decision point”[69], however the details of this process have not been studied in plants.

Most plants fully replicate their genome during endocycles [70], although there are a few exceptions (e.g. various orchid species; [21, 22]). We found very little evidence for over- or under-replication occurring in endocycling maize root cells, unlike the distinctive over- and under-replication found in *Drosophila* endocycles (reviewed in [17] and references therein). Our result is consistent with earlier cytological reports that whole chromosomes, as well as repetitive knobs and centromeres, are completely replicated in the highly endopolyploid maize endosperm [23].

In contrast to the global maintenance of RT, we observed a small fraction of the maize genome that exhibits some difference in RT between the two types of cell cycles. Approximately 11% of the genome showed compensated differences at a stringency level of ≥ 10% difference in replication signal (see Methods). However, with the notable exception of centromeric regions, which are discussed in more detail below, we chose to characterize only the most robust Regions of Altered Timing (RATs), defined by the criteria of containing a core region with compensated differences at a stringency level of ≥ 25% difference in replication signal. These robust non-centromeric RATs comprise only 1.6% of the genome, and the size range of individual regions (39–387 kb, median 138 kb) is consistent with our previous observation that regions of coordinate replication in maize are ∼50–300 kb in size [12]. This may include from one to a few replicons, based on previous estimates of replicon size in monocot plants [71].

The first 1 mm of the maize root contains the meristem and precursors for at least ten different cell types. Only some of these cell types enter the endocycle prior to cell elongation [62]. If there are differences in the RT programs of different cell types, some or all of the non-centromeric RATs may be associated with shifts in the relative contribution of different cell types to the two samples of nuclei, rather than to endocycling *per se*. Research in metazoans has revealed ∼8-20% of their genomes can shift RT between cell types [13, 25, 26, 72–74]. In mammals, these shifts generally involve large regions or “domains” in the megabase size range (reviewed in [16]). These RT domains are much larger than the non-centromeric RATs in maize, even though the maize genome is similar in size to the human and mouse genomes. However, in the much smaller *Drosophila* genome, regions that show timing shifts between cell types are more similar in size to the maize non-centromeric RATs [72, 74].

The vast majority of the non-centromeric RATs involved RT shifts from Earlier-to-Later, with a significant enrichment for not only genes, but genes expressed in the root tip. This result suggests the possibility that RT shifts may be related to shifts in gene expression. Unfortunately, we have been unable to follow transcriptional changes in endocycling nuclei directly, as we have as yet been unable to isolate RNA of sufficient quality to characterize transcripts from fixed nuclei. However, our analysis of activating and repressive histone modifications uncovered only minor changes in the enrichment and location of these marks within RAT genes after endocycle replication. The lack of notable changes in the proportion of RAT genes bearing H3K56ac and H3K4me3 modifications after the endocycle suggests that these histone marks are permissive to changes in RT. Nonetheless, the direction of the change in H3K4me3 enrichment on genes in Earlier-to-Later RATs after endocycle replication (S8B Fig) is consistent with the hypothesis that a shift to later RT may accompany a decrease in gene expression. Many studies have identified a correlation between RT and transcriptional activity (reviewed in [16]), but there are also multiple examples of these processes being uncoupled (e.g. [27, 75]).

In the case of centromeres, it is easy to imagine that the large shifts to later replication are related specifically to endocycling, because endocycling cells no longer require functional centromeres. Though often broken by unmappable and multi-mapping (“blacklist”) regions in the genome assembly, when combined, centromeric RATs are much larger in size than the non-centromeric RATs and cover the majority of each of the seven complex centromeres (S5 Table). These seven centromeres, which are well assembled in the maize B73 RefGen_v4 genome, contain satellite repeats interspersed with retrotransposons [38, 47], enabling almost 50% of our sequencing reads that map to these centromeres to be uniquely positioned. In most species, in which centromeres contain large numbers of tandemly arrayed satellite repeats, it is difficult to map centromeric sequence reads to unique positions and, thus, to fully assess centromeric RT patterns [76]. Though yeast centromeres replicate in early S phase [77–80], most higher eukaryotes replicate centromeres asynchronously through mid to late S phase [54, 81–86]. Many of the reports in higher eukaryotes are based on cytological observations, membrane hybridization, or PCR data with limited resolution. Even a recent genomic analysis of centromeric RT in human cell lines was significantly limited by the quality of the human centromere assemblies, and could only uniquely map ∼15% of centromeric reads [76]. Centromere replication in plant species, assessed mostly by cytological methods, has variously been reported to occur in early, mid or late S [87–90], though it is often unclear if the analysis was of sufficient resolution to distinguish the RT of centromeres from that of adjacent pericentromeres. In contrast, we have provided a high-resolution analysis of the distribution of replication times across maize centromeres, and compared RT of centromeres to adjacent pericentromeres. These analyses revealed several features shared by the RT programs of the seven complex maize centromeres. For example, in mitotic cells there are a few distinct peaks of early replication (e.g. arrowheads in Figs 3 and S12), interspersed with mainly mid replication activity that transitions to late replication at the edges of the functional centromere. In the endocycle, entire centromeres – including regions with early and mid replication activity and the genes, retroelements and *CentC* repeats within them – undergo a shift to later replication. As a result, the RT of the complex centromeres in the endocycle becomes much more similar to that of the immediately adjacent pericentromeric regions, which replicate primarily in late S phase in both mitotic and endocycling cells.

The presence of distinct peaks of early replication in or adjacent to functional centromeres (arrowheads in Fig 3 and S12) is noteworthy because they signify a population preference for initiation in early S phase at these loci. This observation is of particular interest because yeast centromeres contain a replication origin that is the first to initiate on its respective chromosome and plays a role in centromere specification [80]. In maize, there is no evidence that these early regions in centromeres are the first to replicate on the entire chromosome, but they are earlier replicating than their surroundings. Origin mapping experiments (e.g. [91, 92]) would be required to distinguish whether these early regions contain single or small clusters of origins, and the location of any other origins in centromeres that may fire in mid or late S phase.

Unlike complex centromeres, the three simple centromeres of maize show less drastic timing changes, that occur over smaller regions. These simple centromeres are not as well assembled as the complex centromeres [40, 47], and we cannot assess RT for the possibly large portions of these centromeres not present in the genome assembly. One potential interpretation of our results is that the simple centromeres have distinct RT programs that show less timing shift in the endocycle, possibly related to their different sequence composition. Alternatively, the missing portions of the simple centromere assemblies could be replicating more like the complex centromeres. Because simple centromeres are known to primarily contain large *CentC* arrays [40, 47], the second hypothesis is supported by our analysis of mapped *CentC* satellite repeats in all centromeres, which showed that, as a group, these repeats consistently shift RT from mid to late. Another piece of evidence comes from our analysis of complex centromeres, which showed that the magnitude of the RT change tapers off toward the outer edges of the functional centromere. One can speculate that the simple centromere assemblies are comprised mostly of the sequences at the edges of the actual centromere, which would still be anchored to nonrepetitive regions in the genome assembly. As in complex centromeres, these edge sequences might have a smaller RT shift than internal sequences. Future cytological experiments, using a combination of flow sorted EdU-labeled nuclei and techniques for identifying maize chromosomes [93, 94] could help address questions related to the RT of simple centromeres.

The centromere-specific histone variant, CENH3 (also called CENP-A in animal systems) plays an important role in recruiting kinetochore proteins [44–46]. In metazoans, it has been shown that CENP-A is distributed among sister centromeres during replication, but the full complement of new molecules is not redeposited until later [55, 95]. However, there are differences in the timing of deposition of CENH3/CENP-A among eukaryotes. Deposition occurs from S phase to G2 in yeasts, while in plants and protozoans it occurs from late G2 to metaphase, and in metazoans it occurs mostly during G1 (with the exception of some *Drosophila* cell types in metaphase to G1; reviewed in [46, 56, 60]). These interesting differences between phylogenetic groups in the timing of CENH3/CENP-A deposition suggest there may also be differences in the mechanisms and regulation of deposition that need to be explored further [59]. In our analysis of CENH3 enrichment relative to DNA content in maize root cells, the population of 4C nuclei appear to have a full complement of CENH3, which would be consistent with the previous results for plant species. This result suggests a model in which the sub-population of 4C cells entering the endocycle also carry a full complement of CENH3. If that model is correct, our data for 8C nuclei imply that CENH3 is only partially replaced after DNA replication in the endocycle. Because the population of 8C nuclei we analyzed likely represents a mixture of cells that recently exited endocycle S phase and others that exited some time ago we cannot determine whether CENH3 will be fully restored in all cells at a later time. However, it is clear that the ratio of CENH3 to DNA is not immediately restored, and the lower ratio is widely distributed across all ten centromeres.

It is unlikely that endocycling cells will ever re-enter the mitotic cycle [1, 96, 97], and it is not clear why endocycling cells would maintain or redeposit CENH3 nucleosomes at all unless CENH3 has roles outside of mitotic cell division. A recent study in *Drosophila* midgut cells found that CENP-A is required even in post-mitotic and differentiated cells, and proposed that the loading of CENP-A in endocycling cells is essential for maintaining chromosome cohesion [98]. This possibility has not yet been tested.

Centromeres are considered to be epigenetically specified, as there are no unique sequences in the functional centromere that are not also found in the adjacent pericentromere (e.g. reviewed in [44, 99, 100]). With this in mind, we tested whether changes in enrichment levels of CENH3 nucleosomes, or several modifications to canonical H3 nucleosomes, could explain the large shift to later replication of centromeres in endocycling cells. These studies only uncovered very small changes in activating and repressive histone H3 modifications in centromeres after endocycle replication. The magnitude of the change in CENH3, while somewhat larger, was not on the scale of the change in RT. It is possible that the decrease in dosage of CENH3 proteins has an effect on the recruitment of replication proteins, as has been proposed in the yeast *Candida albicans* [80]. If replication proteins were not recruited as efficiently, this could contribute to a delay in replication time of the centromere. It is also possible that more significant changes might be found in epigenetic marks that we did not investigate, for example changes in DNA methylation patterns or other histone post-translational modifications. A variety of modifications to CENP-A nucleosomes have been identified, (reviewed in [101]), but very little is known about CENH3 modifications in plants [102, 103], highlighting an area for future research. Experiments in human cells identified cell cycle related interchanges of acetylation, monomethylation and ubiquitination at the lysine 124 residue of CENP-A [104, 105]. Mutations of this residue led to replication defects and alterations to centromeric RT [105]. Another interesting question is whether changes in chromatin conformation or 3D positioning in the nucleus are associated with the large shift in centromeric RT. In mammals, RT is considered a functional readout of large-scale chromatin structure [16, 27, 73], and regions that shift RT have been shown to also change 3D localization [106]. Additionally, a study in mouse showed that when late replicating pericentric heterochromatin was experimentally repositioned to the nuclear periphery, a location where mid replicating chromatin is usually found in that system, the RT of those regions was advanced [107].

Investigating the interplay of chromatin environment, gene transcription and DNA replication in plant systems, particularly in important crop species, has proven difficult in the past. Numerous reasons for these difficulties exist, for example, plants have cell walls and are rich in nucleases, actively dividing cells are sequestered in tiny meristematic regions, and many genomes have a high content of retrotransposons and other repeats. As a result, understanding of such critical areas has lagged behind that in yeast and animal systems. However, with recent progress in assembling genomic resources and anticipated advances in the ability to isolate individual cell types [108], perform sophisticated analyses of genome conformation [109, 110] and follow individual chromosome regions using elegant cytological paints [94], the maize root tip system is poised to contribute to rapid progress in these and many other important areas of plant genome biology.

## METHODS

### Plant material

Seeds of *Zea mays* inbred line B73 (GRIN NPGS PI 550473) were germinated on damp paper towels and grown for three days. Seedling roots were labeled by immersion in sterile water containing 25 μM EdU (Life Technologies) for 20 min, using growth and experimental conditions described previously [8, 9, 12]. Biological replicate material was grown independently and harvested on different days. For the endocycle Repli-seq experiment, after rinsing roots well with sterile water, the 1–3 mm segments (Fig 1A) were excised from primary and seminal roots. The root segments were fixed, washed and snap-frozen as described previously [9].

### Flow cytometry and sorting of root nuclei

Details of the flow sorting for Repli-seq analysis were described previously [9, 12]. Briefly, nuclei were isolated from the fixed root segments, and the incorporated EdU was conjugated to AF-488 using a Click-iT® EdU Alexa Fluor 488 Imaging Kit (Life Technologies). The nuclei were then resuspended in cell lysis buffer (CLB) [9] containing 2 μg/mL DAPI and 40 μg/mL Ribonuclease A and filtered through a CellTrics® 20-μm nylon mesh filter (Partec) just before flow sorting on an InFlux™ flow cytometer (BD Biosciences) equipped with UV (355 nm) and blue (488 nm) lasers. Nuclei prepared from the 1–3 mm root segments were sorted to collect populations of EdU/AF-488-labeled nuclei with DNA contents in three defined sub-stage gates between 4C and 8C, corresponding to early, mid and late S phase of the endocycle. The early endocycle gate was shifted slightly to the right to exclude mitotic nuclei in late S phase (Fig 1C). For each biological replicate, between 50,000 and 200,000 nuclei were sorted from each fraction of the endocycle S phase. A small sample of nuclei from each gate was sorted into CLB buffer containing DAPI and reanalyzed to determine the sort purity (S1 Fig). Sorting and reanalysis details for the mitotic nuclei are described in [12].

For ChIP-seq experiments, roots were labeled with EdU, and nuclei were isolated from 0–3 mm (H3K27me3 and H3K4me3) or 0–5 mm (H3K56ac) root segments and conjugated to AF-488 as described above. The 2C, 4C and 8C unlabeled, non S-phase populations of nuclei were sorted into 2× extraction buffer 2 (EB2) [111] using the same sorting conditions as in Wear et al. [12]. After sorting, the 2× EB2 was diluted to 1× with 1× STE. All flow cytometry data were analyzed using FlowJo v10.0.6 (TreeStar, Inc.) as described in Wear et al. [12].

### DNA and chromatin immunoprecipitations

For endocycle Repli-seq samples, reversal of formaldehyde cross links, nuclear DNA purification and isolation, DNA shearing, EdU/AF-488 DNA immunoprecipitation with an anti-Alexa Fluor 488 antibody (Molecular Probes, #A-11094, lot 895897), and DNA fragment purification were performed as described in Wear et al. [12].

ChIP procedures were performed as in Wear et al. [12] except the chromatin was sheared using a Covaris S220 ultrasonicator to an average fragment size of 200 bp using a peak incident power of 140 W, 10% duty cycle, and 200 cycles per burst for 6 min. Three percent of the chromatin volume was set aside to use as the input control for each of the 2C, 4C and 8C samples and frozen at −70°C until the formaldehyde cross link reversal step. The antibodies used for ChIP were as follows: *Zea mays* anti-CENH3 antibody at a 1:250 dilution (gift from R.K. Dawe) [48], anti-H3K9me2 antibody at a 1:25 dilution (Cell Signaling Technologies; 9753, lot 4), anti-H3K56ac antibody at a 1:200 dilution (Millipore; 07-677, lot DAM1462569), anti-H3K4me3 antibody at a 1:300 dilution (Millipore; 07-473, lot DAM1779237) and anti-H3K27me3 antibody at a 1:300 dilution (Millipore; 07-449, lot 2,275,589). See S18 Fig for antibody validation experiments for anti-H3K9me2 and anti-CENH3.

### Library construction and sequencing

For Repli-seq and ChIP-seq samples, the final purified DNA was used to construct paired-end libraries as described [12]. After adapter ligation, all samples underwent 17 cycles of PCR. For each Repli-seq or ChIP-seq experiment, individual samples from three biological replicates collected on different days were barcoded, pooled and sequenced on either the Illumina HiSeq 2000 or NextSeq platforms. However, in the case of the Repli-seq mitotic late-S samples and CENH3 ChIP 4C samples, one biological replicate failed during library generation or sequencing, resulting in data from only two biological replicates. Repli-seq and ChIP-seq read mapping statistics are shown in S1 Spreadsheet.

### Replication timing data analysis

Trimming and quality control of 100-bp paired-end Repli-seq reads were carried out as described previously [12], and reads were aligned to the maize B73 RefGen_v4 reference genome [38] (Ensembl Plants release 33; ftp://ftp.ensemblgenomes.org/pub/plants/release-33/gff3/zea_mays/) using BWA-MEM v0.7.12 with default parameters [112]. Redundant reads resulting from PCR amplification were removed from each of the alignment files using Picard (http://broadinstitute.github.io/picard/) and SAMtools [113]. Properly paired, uniquely mapping reads (MAPQ score > 10) were retained with SAMtools [113] for downstream analysis. The resulting mitotic Repli-seq data were more than 3× the sequencing coverage of the endocycle Repli-seq data (S1 Spreadsheet). Repli-seq results are robust at various sequencing depths [14], but to ensure that the mitotic and endocycle data were comparable, the reads were downsampled by a uniform random process using a custom python script incorporating the BEDTools suite [114] to a total of 65.7 million reads per sample, representing almost 3× genome coverage for each S-phase fraction (S1 Spreadsheet). We preferred this to normalization so that any possible sampling bias due to sequencing depth would be similar in all samples.

Repli-seq data were analyzed using *Repliscan* [14]. Individual biological replicates of Repli-seq data were independently analyzed, and after finding good correlation between replicates (Pearson correlation coefficients from 0.80–0.99; S4 Fig) the replicates were aggregated by sum and normalized to 1× genome coverage using the reads per genomic content (RPGC) method. The following changes from the *Repliscan* default parameters described in [12] were used. Read densities were aggregated in 3-kb windows across the genome (parameter *-w* 3000). Additionally, we customized the cutoff for reducing type one errors which excluded genomic windows with extremely low coverage in the 2C reference sample. To identify these low read mapping windows, which we labeled “blacklist”, *Repliscan* log-transformed the read counts from the pre-replicative 2C reference sample and windows with read counts in the lower 2.5% tail of a fitted normal distribution were excluded from all samples (parameter *--pcut* 2.5-100). The upper 2.5% tail containing extremely high coverage windows or “spikes” was not removed at this step, because we found that these data spikes were adequately normalized in the subsequent step of dividing each 3-kb window in the S-phase samples by the 2C reference data – which also normalized for sequencing biases and collapsed repeats (S3 Fig). The data were then Haar wavelet smoothed [14] to produce the final profiles for early, mid and late S-phase replication signals in the mitotic cycle and endocycle. Processed data files, formatted for the Integrative Genomics Viewer (IGV) [115], are available for download from CyVerse (formerly the iPlant Collaborative; [116]) via the information in S1 Spreadsheet.

### Identifying regions of altered replication timing

The difference between normalized signal profiles of mitotic and endocycle Repli-seq data for early, mid, and late S was calculated in 3-kb windows, and the maximum negative and positive differences were then calculated for each chromosome and averaged. Regions showing a timing difference of ≥ 25% (difference in replication signal ≥ 1.0) or ≥ 10% (difference in replication signal ≥ 0.4) of the total range of differences in each profile were identified (S1 Table; S5 Fig) using the data filter tool in SAS JMP Pro v14 (SAS Institute Inc.). Windows were kept in the analysis only if their timing differences were “compensated” by opposite timing difference(s) of ≥ 25% or ≥ 10%, respectively, in one or both of the other two S-phase fractions. For example, a decrease in early replication signal in endocycling cells must be compensated by an increase in mid and/or late S-phase signal in the same cell population. Adjacent 3-kb windows with timing differences that met either the ≥ 10% or ≥ 25% threshold were merged, keeping the two files separate, using mergeBED in the BEDTools suite, and allowing a 6 kb gap distance (parameter *- d* 6000) [114]. This initial step resulted in many very small regions being identified (S2 Table). As a second step, if ≥ 10% regions were immediately adjacent to ≥ 25% regions, they were merged together using mergeBED to highlight larger regions of contiguous change (S2 Table). Only regions that contained at least one ≥ 25% region were kept for further analysis, and termed regions of alternate timing (RATs). By requiring a ≥ 25% RT change core region to be included, all of the stand-alone, extremely small regions (< 24 kb) were effectively filtered out, without the requirement of an arbitrary size filter. RATs were categorized into three groups: 1) later in mitotic to earlier in endocycle (Later-to-Earlier), 2) earlier in mitotic to later in endocycle (Earlier-to-Later) and 3) a subset of the Earlier-to-Later RATS that were located in the previously identified functional centromeres (Earlier-to-Later-CEN) (coordinates from [38]). There were no Later-to-Earlier-CEN RATs. For a list of RAT regions, including genomic coordinates and genes within them, see S2 and S3 Spreadsheets.

### ChIP-seq data analysis

ChIP-seq reads for H3K27me3, H3K4me3, H3K56ac (100-bp paired-end reads), H3K9me2 and CENH3 (150-bp paired-end reads) were trimmed, mapped to maize B73 RefGen_v4.33, and filtered to retain only properly-paired, uniquely-mapped reads (MAPQ score > 10) as described above for Repli-seq reads. The 2C ChIP and input data for H3K27me3, H3K4me3, H3K56ac is from [12], while the 4C and 8C ChIP data was generated for this study, see S1 Spreadsheet. For details on peak calling and analysis for H3K27me3, H3K4me3, H3K56ac, see S1 Text.

For visualization of CENH3 localization in 2C, 4C and 8C nuclei, read counts for individual biological replicates of CENH3 or input samples were scaled to 1× genome coverage using the reads per genomic content (RPGC) method. Biological replicate data had good agreement (Pearson’s correlation coefficient values between biological replicates of 0.97-0.99; S1 Spreadsheet), and were merged and scaled again to 1× coverage so the samples would be comparable. CENH3 scaled read counts in each 3-kb window were divided by the scaled read counts from the input sample for the corresponding ploidy level, resulting in CENH3 fold enrichment values relative to input.

To compare CENH3 enrichment relative to DNA content in 2C, 4C and 8C cells over entire centromeres, we calculated the percent of total CENH3 reads found in a given centromere (using coordinates from [38]), divided by the percent of total input reads corresponding to that centromere. This was done separately for individual biological replicates; we then calculated the mean fold enrichment estimates. H3K9me2 fold enrichment over entire centromeres and pericentromeres was calculated in the same way.

### Genomic features

The maize filtered gene set Zm00001d.2 annotation from B73 RefGen_v4 [38] was downloaded from Ensembl Plants (ftp://ftp.ensemblgenomes.org/pub/plants/release-33/gff3/zea_mays/). The updated B73 Refgen_v4 TEv2 disjoined annotation [39] was downloaded from http://mcstitzer.github.io/maize_TEs. Coordinates for mapped *CentC* satellite repeat regions are described in Gent et al. [40]. The percent AT content was calculated in 3-kb static windows across the genome.

### Analysis of features in RATs and random permutation analysis

We tested the association of various genomic features with the non-CEN RAT categories by determining the overlap of a particular feature with each RAT type. The coordinates for genomic features (genes, expressed genes, TE superfamilies) were intersected with RAT coordinate intervals using intersectBED (parameters *-wa -wb*) in the BEDtools suite [114]. The percent of RATs containing a feature or the percent coverage of genes and TE superfamilies were computed and compared to values for the genome as a whole. The number of genes per RAT was also determined using intersectBED (parameter *-u*).

For comparison, the coordinates for the non-CEN Earlier-to-Later and Later-to-Earlier RAT sets were randomly shuffled around the genome, excluding functional centromeres, using BEDTools shuffle [114]. These random sets preserved the number of regions and region size of the original RAT sets, and are labeled “EtoL shuffle1” and “LtoE shuffle1” for the Earlier-to-Later and Later-to-Earlier RATs, respectively. When there appeared to be differences in the observed overlap values with genomic features between non-CEN RATs and their corresponding random shuffle sets, a permutation or feature randomization test, as described in [12] was used to assess the statistical significance of the observed value. To do so, the coordinates for the non-CEN RAT sets were randomly shuffled around the genome 1000 times, as described above.

### Analysis of features in centromeres and pericentromeres

For comparison to CEN regions (coordinates from [38]), pericentromeres were arbitrarily defined as the ± 1 Mb flanking each CEN. In the case of chromosome 9, the pericentromere included the ± 1 Mb flanking both CEN 9a and 9b. Replication timing signal values in CENs and pericentromeres were intersected with genes, *CRM1* and *CRM2* families and mapped *CentC* regions using intersectBED (parameters *-wa -wb*) in the BEDtools suite [114]. Only elements that covered at least half of a 3-kb window of Repli-seq data were included in Fig 4, while elements with any amount of overlap were included in S14 Fig. Additionally, if a single gene or *CRM* element spanned more than one of the 3-kb windows, the replication signals were averaged using mergeBED (parameter *-o* mean) to compute a single value for the entire gene or element.

## ACKNOWLEDGEMENTS

We thank the following people for helpful discussions of this work: Ashley Brooks, Emily Wheeler and Hank Bass. We are grateful to Kelly Dawe and Jonathan Gent for sharing the maize CENH3 antibody, mapped *CentC* data and advice. We thank Patrick Mulvaney and James Harrison Priode for help with harvesting plant material.

## SUPPORTING INFORMATION

### S1 Text. Supplemental Methods

**S1 Fig. (related to Fig 1) Assessment of purity of flow sorted endocycling nuclei.** Maize root tip nuclei were isolated from the 1–3 mm root region and sorted on a BD InFlux flow sorter. A small sample from each of the three S-phase sort gates was re-analyzed to determine the purity of the sorted nuclei. Histograms of relative DNA content (DAPI fluorescence) from re-analyzed sorted nuclei are overlaid for early (E), mid (M), and late (L) S-phase gates from the endocycle arc to show the separation between sorted samples. Similar separation was found for sorted early, mid and late nuclei from the mitotic cycle (see Supplemental Fig. 1 in [12]). The histogram of relative DNA content for the entire unsorted nuclei population (black line) is shown for reference.

**S2 Fig. (related to Fig 1) Genomic copy number analysis.** Whole genome sequence data from sorted non S-phase 2C, 4C and 8C nuclei were used to assess copy number per DNA content across the genome. To better represent the copy number of repeat regions, the primary alignment location for each read pair – even those that map to multiple locations – were included in the analysis. (**A and B**) Histograms of the normalized read frequency ratios, calculated in 5-kb static windows, for 2C/4C (**A**) and 8C/4C (**B**) nuclei. The black dashed lines indicate the overall mean and the red dashed lines indicate ± 2 S. D. from the mean. (**C**) The 8C/4C read frequency ratios plotted as a function of genomic location, which shows that the values outside ± 2 S. D. all occur as singleton 5-kb windows. (**D and E**) We used consensus sequences for 45S rDNA and *knob180* (**D**), and for 5S rDNA, *TR-1*, *CentC* and *CRM1–4* families (**E**) to individually query all of the trimmed whole genome sequence reads using BLAST software and a non-stringent E value to allow for variants of each repeat (S1 Text). The mean percentage of total reads that align to each repeat type was calculated for three biological replicates of 2C, 4C and 8C data. Black dots represent the individual biological replicate values. The apparent slight under-replication of several elements (e.g., *knob180* and *CRM2*) is not statistically significant.

**S3 Fig. (related to Figs 1 and 3) Example of Repli-seq data processing with *Repliscan*.** An example region from CEN 10 is shown to illustrate that the pre-replicative 2C reference data effectively normalizes spikes of signal in the S-phase data. (**A and B**) Read densities were calculated in 3-kb windows for the 2C reference (**A**) and each S-phase sample (endocycle late profile shown; **B**). After excluding blacklist regions (e.g. unmappable and multi-mapping regions), reads were scaled for overall sequence depth in each sample. (**C**) Scaled reads in each S-phase sample were normalized by making a ratio to 2C reference scaled reads in each 3-kb window. (**D**) Replication signal profiles were smoothed using a Haar wavelet transform to remove noise without altering peak boundaries.

**S4 Fig. (related to Fig 1) Pearson’s correlation coefficient values between individual biological replicates of mitotic and endocycle Repli-seq data.** (**A and B**) Biological replicates (BR) of early (E), mid (M) and late (L) Repli-seq data for the mitotic cycle (Mit; panel **A**) and endocycle (En; panel **B**) was analyzed independently using *Repliscan* [14]. The agreement between biological replicates was assessed by calculating Pearson’s correlation coefficients. (**C**) The Pearson’s correlation coefficients for E, M, L data between mitotic cycle and endocycle.

**S5 Fig. (related to Fig 2) Boxplots of differences in early, mid and late replication signal profiles for each chromosome.** Differences in replication (dRT) signal were calculated by subtracting the mitotic signal from the endocycle signal for early (E), mid (M) and late (L) S-phase fractions in each 3-kb window across the genome. The distributions of dRT signal values are represented as violin plots for each chromosome. Median values are indicated by colored squares and 1.5 x IQR of the distribution is indicated by colored whisker lines. Dashed lines indicate the thresholds used in subsequent steps for identifying RATs (≥ 10% and ≥ 25% of the total difference range; S1 Table).

**S6 Fig. (related to Fig 2) Additional examples of non-CEN RATs.** (**A–F**) Example regions on chromosomes 1 (**A**), 3 (**B**), 4 (**C**), 5 (**D**), 6 (**E**) and 7 (**F**) that include RATs. See main text Fig 2 legend for description. Dashed boxes denote regions with some level of RT difference in which the magnitude of the difference did not meet our ≥ 25% criterion (boxes labeled “a” in panels **A**, **B**, **C** and **F**), or in which the change in one S-phase fraction was not compensated by an opposite change in at least one other S-phase fraction (boxes labeled “b” in panels **C** and **D**).

**S7 Fig. (related to Fig 2 and** Table 1**) Permutation analysis of the percentage overlap of non-CEN RATs and genes.** (**A–D**) The percentage of RATs that overlap genes (**A and B**) or expressed genes (**C and D**) was calculated for non-CEN RATS and 1000 randomly shuffled sets (see Methods). The observed percentage for RATs (red line) and the frequency distribution of the random sets (green) are plotted.

**S8 Fig. (related to Fig 2) Activating and repressive histone marks in non-CEN RATs.** To assess whether changes in selected histone modifications related to gene transcription and chromatin accessibility occur in RATs, ChIP-seq data was generated for H3K56ac and H3K4me3 (active transcription and early replication) and H3K27me (repressive transcription and facultative heterochromatin) from sorted non S-phase 2C, 4C and 8C nuclei. (**A–C**) The distributions of fold enrichment values for H3K56ac (**A**), H3K4me3 (**B**) and H3K27me3 (**C**) peaks in expressed and non-expressed genes (see S1 Text) in 2C, 4C and 8C nuclei are plotted as boxplots for Later-to-Earlier and Earlier-to-Later RATs and their corresponding randomly shuffled sets (see Methods). Asterisks indicate statistically significant differences by the non-parametric Steel-Dwass-Critchlow-Fligner test at the following *P* value levels: ***, *P* < 0.0001; **, *P* < 0.001; *, *P* < 0.01. The increase in the fold enrichment of H3K56ac for expressed genes in Earlier-to-Later RATs (panel **A**) may be associated with increases in peak enrichment we observed near the 3’ end of some genes. (**D**) The count and percentage of expressed and non-expressed genes with each histone modification shown in the boxplots in panels **A–C**. The 8C/2C ratio of genes with each mark is also shown to demonstrate there is very little change in the number of genes with each mark. The total number of expressed and non-expressed genes in each RAT or random category are shown at the bottom for reference.

**S9 Fig. (related to Fig 2) Gene ontology analysis of genes in non-CEN RATs.** Using the Plant GO slim ontology subset, we identified 44 significant GO terms in the biological process (P), molecular function (F), and cellular component (C) GO categories that were enriched in expressed genes (S1 Text; S3 Spreadsheet) in Earlier-to-Later RATs. Genes in the corresponding randomly shuffled set shared a few of the significantly enriched cellular component terms as genes in Earlier-to-Later RATs, suggesting that these terms may be related to common components of the root, and not RATs specifically. The total number of expressed genes in each input gene list was as follows: Later-to-Earlier RATs, 52; LtoE shuffle1 random regions, 68; Earlier-to-Later RATs, 292; EtoL shuffle1 random regions, 275.

**S10 Fig. (related to Fig 2) AT content composition in non-CEN RATs.** (**A**) The distributions of percent AT content, calculated in 3-kb static windows, for Later-to-Earlier and Earlier-to-Later non-CEN RATs and the corresponding random shuffle sets are plotted as boxplots. Values outside the boxplot whiskers (1.5 x IQR) are represented as grey dots. The dashed line indicates the genome wide median value.

**S11 Fig. (related to Fig 3) Uniquely mapping Repli-seq reads in centromeres.** The average percentage of centromeric reads that map to unique locations is shown for each replication timing sample. Black dots represent the individual values for biological replicates.

**S12 Fig. (related to Fig 3) Replication signal profiles and RATs in complex and simple centromeres.** 5-Mb regions are shown for complex CENs 2, 3, 4, 5, and 8 and simple CEN 7. See main text Fig 3 legend for description.

**S13 Fig. (related to Fig 3) Timing differences in centromeres and pericentromeres.** Timing differences (endocycle minus mitotic) between early (**A and D**), mid (**B and E**) and late (**C and F**) profiles for each centromere and corresponding pericentromere (± 1 Mb) were calculated in 100-kb static windows. In panels **D**, **E**, and **F** asterisks indicate difference values from windows where an Earlier-to-Later-CEN RAT extends past the called CEN boundary [38] into the pericentromere; open circles indicate windows that contain a non-CEN Earlier-to-Later RAT that met our compensation criteria.

**S14 Fig. (related to Fig 4) Replication times for all genomic features in complex and simple centromeres and corresponding pericentromeres.** All elements within centromeres and pericentromeres are included, not just those that cover at least half of a 3-kb window, as in Fig 4. See main text Fig 4 legend for description.

**S15 Fig. (related to Fig 4) Activating and repressive histone mark peaks of enrichment in centromeres.** ChIP-seq data were generated for H3K56ac, H3K4me3 (active transcription) and H3K27me (repressive transcription) from 2C, 4C and 8C nuclei. (**A–C**) The fold enrichment values for peaks in expressed and non-expressed genes for H3K56ac (**A**), H3K4me3 (**B**) and H3K27me3 (**C**) in 2C, 4C and 8C nuclei. Red lines indicate the median value. (**D**) The number of expressed and non-expressed genes with each mark in 2C, 4C and 8C nuclei.

**S16 Fig. (related to Fig 5) H3K9me2 fold enrichment relative to DNA content in complex and simple centromeres.** We used the ChIP-seq datasets from 2C, 4C and 8C nuclei to estimate the H3K9me2 average fold enrichment relative to DNA content by calculating the percent of total H3K9me2 reads found in a given centromere (**A and B**) using coordinates from [38] or pericentromere (**C and D**) and dividing by the percent of total input reads corresponding to that centromere or pericentromere. Black dots represent the individual values from biological replicates.

**S17 Fig. (related to Fig 5) CENH3 localization and enrichment in mitotic and endocycling centromeres.** (**A**) CENH3 localization patterns for 2C, 4C and 8C nuclei for CEN 1–CEN 8. (**B**) CENH3 average fold enrichment relative to DNA content for complex and simple centromeres. See main text Fig 5 legend for CEN 9 and CEN 10 localization patterns and description.

**S18 Fig. (related to Fig 5) ChIP-qPCR antibody validations for anti-CENH3 and anti-H3K9me2 antibodies.** The percentage of input (%IP) was calculated for various antibody dilutions and primer sets for the *Zea mays* anti-CENH3 antibody **(A)** and anti-H3K9me2 antibody **(B)**. Black dots in panel **A** represent the individual values from two biological replicates. Positive control primer sets (*CRM2* and Copia retrotransposons) and negative control primer sets (18S rDNA and Actin1 UTR) were used. The no antibody control (NoAB) values are too small to see on the graph. See S1 Text for Supplemental Methods.

**S1 Table. (related to Fig 2) Replication timing signal differences and thresholds.** The difference in replication timing signal between mitotic and endocycle profiles (endocycle minus mitotic) was calculated for each 3-kb window across the genome. The maximum negative difference value, which indicates a higher signal in the mitotic cycle, and the maximum positive difference value, which indicates a higher signal in the endocycle, are shown for early and late profiles. The average total difference range between these two values was used to calculate percentage thresholds for identifying RATs (see S2 Table and main text).

**S2 Table. (related to Fig 2) Summary statistics of preliminary RAT calling steps.** The thresholds from S1 Table (≥ 10% or ≥ 25%) were used to identify regions with RT difference in early or late S phase that were compensated by difference(s) with an opposite sign in one or both of the other two S-phase fractions (early + mid or mid + late) with greater than or equal to the same magnitude. The count, minimum, maximum and median region size, and the total coverage of the B73 RefGen_v4 genome are shown. Final robust RATs included at least one core region with a ≥ 25% RT difference, but immediately adjacent regions of ≥ 10% differences were merged together with the ≥ 25% regions to identify larger regions of contiguous change.

**S3 Table. (related to Fig 2) Gene summary in non-CEN RATs.** The percent of RATs that contain genes, the total number of genes and expressed genes and the mean gene count per RAT are shown.

**S4 Table. (related to Fig 3) Permutation analysis results for gene and TE coverage in non-CEN RATs.** The permutation *P* values derived from calculating percent coverage in 1000 random permutations of each RAT set (e.g. see S7 Fig). All permutation *P* values shown are associated with a test for whether the observed percent coverage value is greater than expected by chance, unless marked “NEG” which indicates the *P* value is associated with a test for whether the observed percent coverage value is less than expected by chance.

**S5 Table. (related to Fig 3) Cumulative RAT coverage in centromeres.** The cumulative coverage and number of RATs called in each centromere are shown. For reference, the previously determined centromere sizes are shown [38], as well as the sizes after unmappable regions are subtracted out. There are also some unmappable regions of unknown size missing from the genome assembly [38], which we cannot account for here.

**S6 Table. (related to Fig 4) Compensated timing shifts in complex centromeres and corresponding pericentromeres.** We calculated the total number of 3-kb windows in complex centromeres and pericentromeres (± 1 Mb), as well as the number of windows that show timing shifts that are compensated (threshold ≥ 10%) by equal and opposite shifts in the other two S-phase fractions.

**S7 Table. (related to Fig 5) CENH3 average fold enrichment relative to DNA content in centromeres.** CENH3 fold enrichment relative to DNA content and the ratio of enrichments between 4C and 2C and 8C and 4C are shown for each centromere. Fold enrichment values are the mean ± S. D. of three biological replicates for 2C and 8C and two biological replicates of 4C. See main text Fig 5 legend for further description. Two sets of theoretical ratio values are also presented. The first set, labeled “proportional redeposition”, corresponds to the hypothesis that CENH3 is diluted relative to total DNA during replication, and is then redeposited to a level proportional to the DNA content during the subsequent gap phase. The second set, labeled “no redeposition”, corresponds to an alternate hypothesis that CENH3 is diluted relative to total DNA during replication, and is not redeposited in the subsequent gap phase.

**S1 Spreadsheet. (related to Figs 1–5) Mapping statistics and data availability for all included datasets.**

**S2 Spreadsheet. (related to Figs 2 and 3) RAT regions list.**

**S3 Spreadsheet. (related to Figs 2 and 3) Genes found in RATs.**

