## Supplemental Figures for "Comparing DNA replication programs reveals large timing shifts at centromeres of endocycling cells in maize roots"

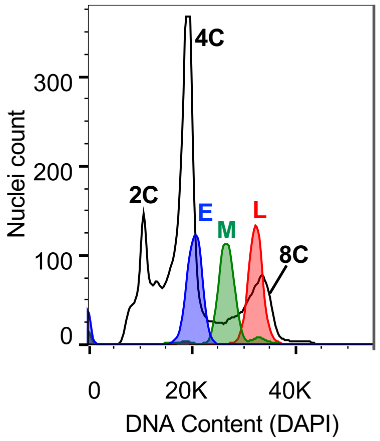

4C
2C
E
M
L
8C
300
200
100
0
Nuclei count
0
20K
40K
DNA Content (DAPI)

S2 Fig.

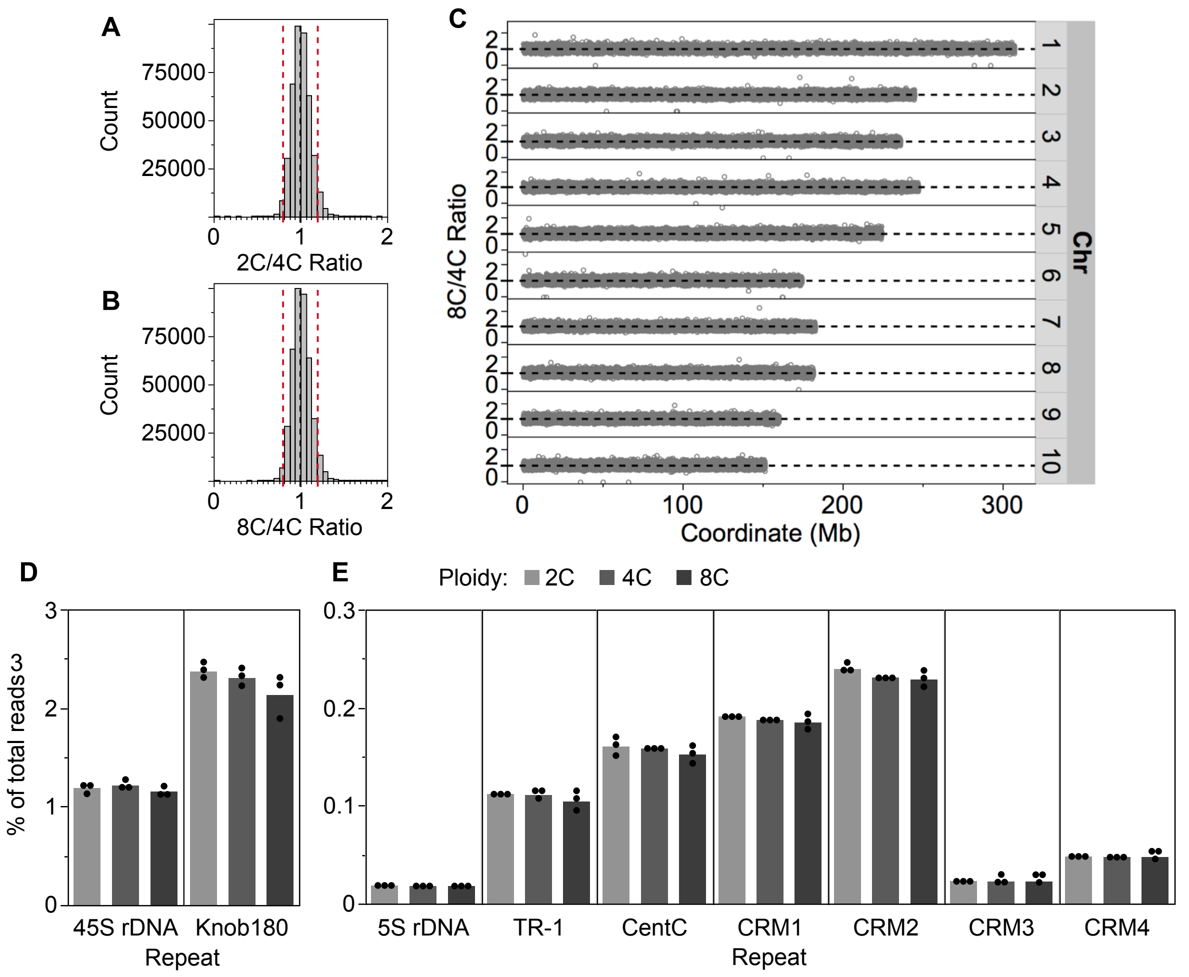
A
Count
75000
50000
25000
0
1
2
2C/4C Ratio
B
Count
75000
50000
25000
0
1
2
8C/4C Ratio
C
8C/4C Ratio
2
0
1
2
3
4
5
6
7
8
9
10
Chr
0
100
200
300
Coordinate (Mb)
D
% of total reads
3
2
1
0
45S rDNA
Knob180
Repeat
E
Ploidy:
2C
4C
8C
0.3
0.2
0.1
0
5S rDNA
TR-1
CentC
CRM1
CRM2
CRM3
CRM4
Repeat

S3 Fig.

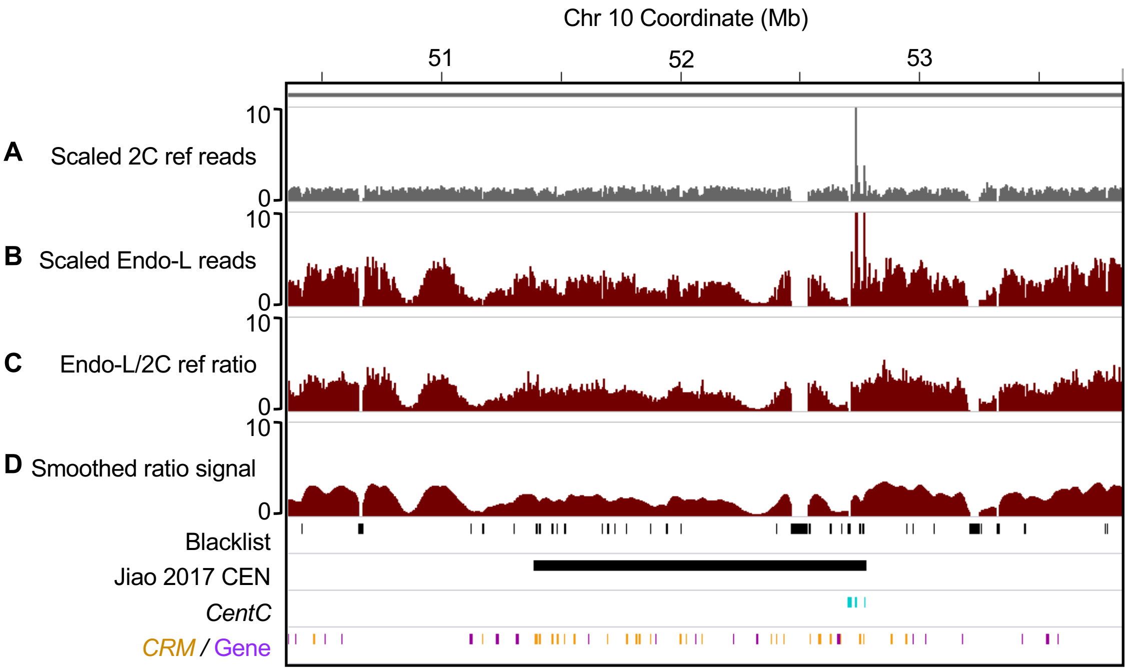

Chr 10 Coordinate (Mb)
51
52
53
10
0
A Scaled 2C ref reads
B Scaled Endo-L reads
C Endo-L/2C ref ratio
D Smoothed ratio signal
Blacklist
Jiao 2017 CEN
CentC
CRM / Gene

S4 Fig.

A

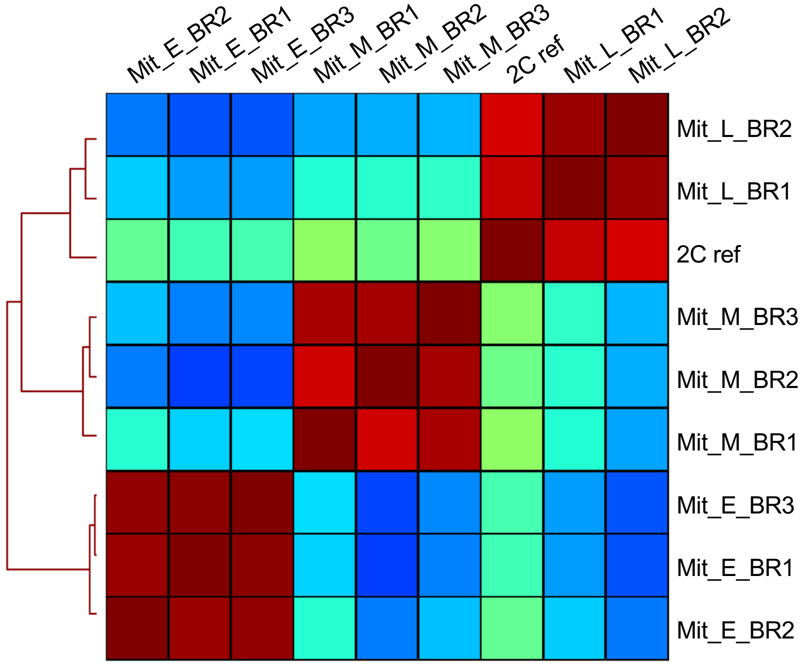

B

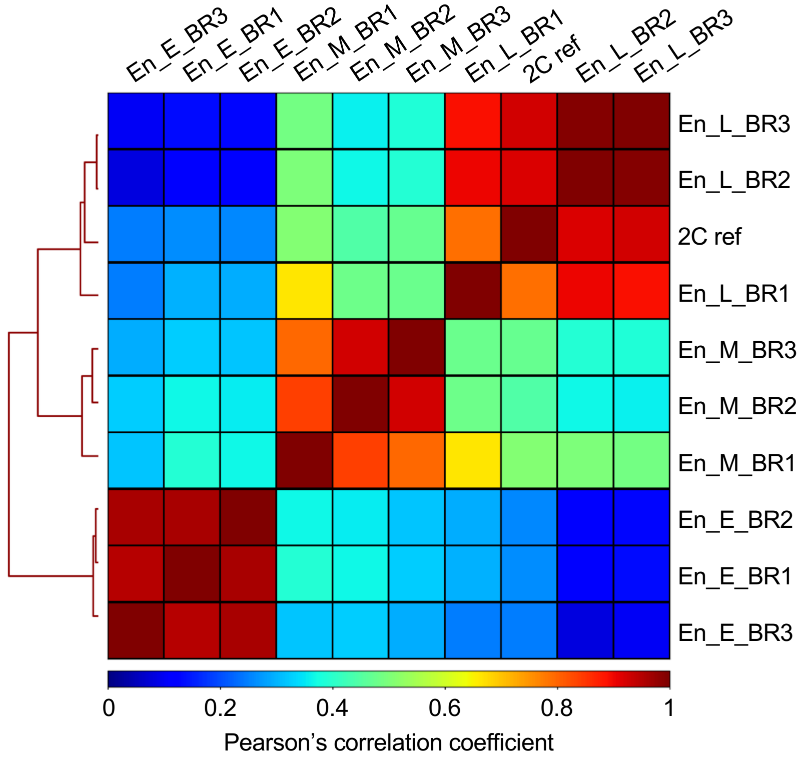

C

| Mitotic vs. Endocycle correlation coefficients | |
|---|---|
| E | 0.91 |
| M | 0.89 |
| L | 0.96 |

S5 Fig.

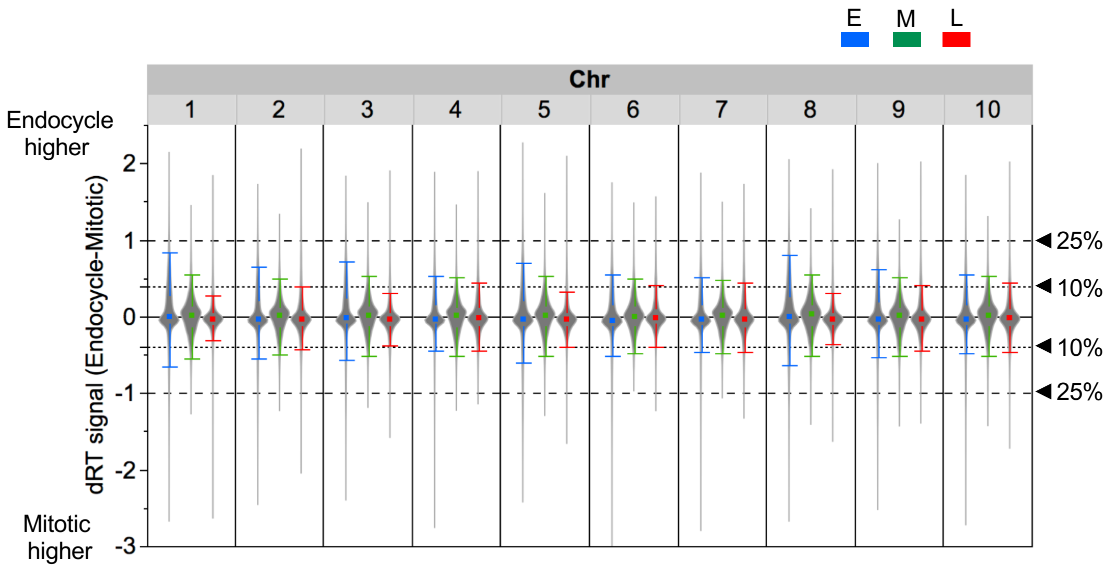
E
M
L
Chr
1
2
3
4
5
6
7
8
9
10
Endocycle higher
Mitotic higher
dRT signal (Endocycle-Mitotic)
2
1
0
-1
-2
-3
25%
10%
10%
25%

S6 Fig.

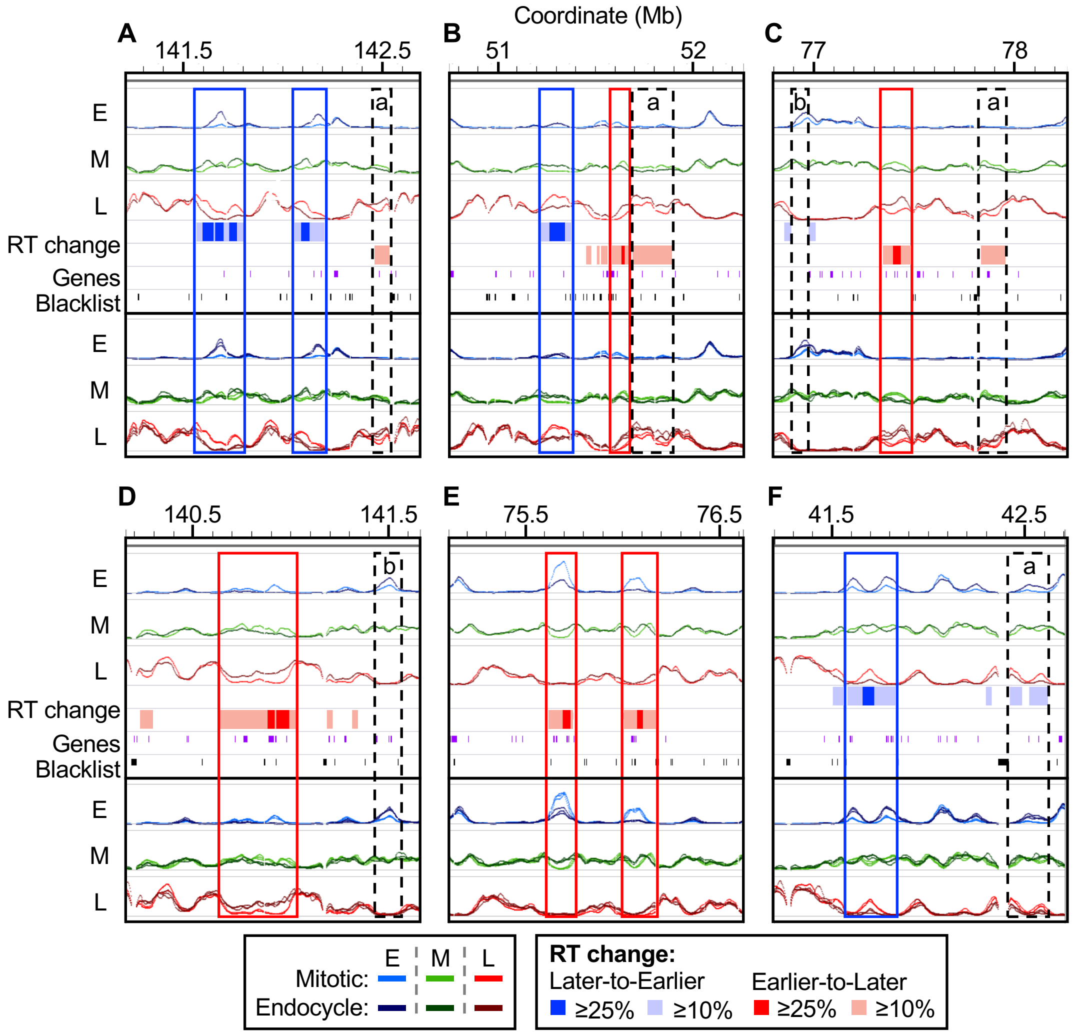
Coordinate (Mb)
A
141.5
142.5
B
51
52
C
77
78
a
b
E
M
L
RT change
Genes
Blacklist
D
140.5
141.5
E
75.5
76.5
F
41.5
42.5
Mitotic:
Endocycle:
RT change:
Later-to-Earlier
Earlier-to-Later
≥25%
≥10%

S7 Fig.

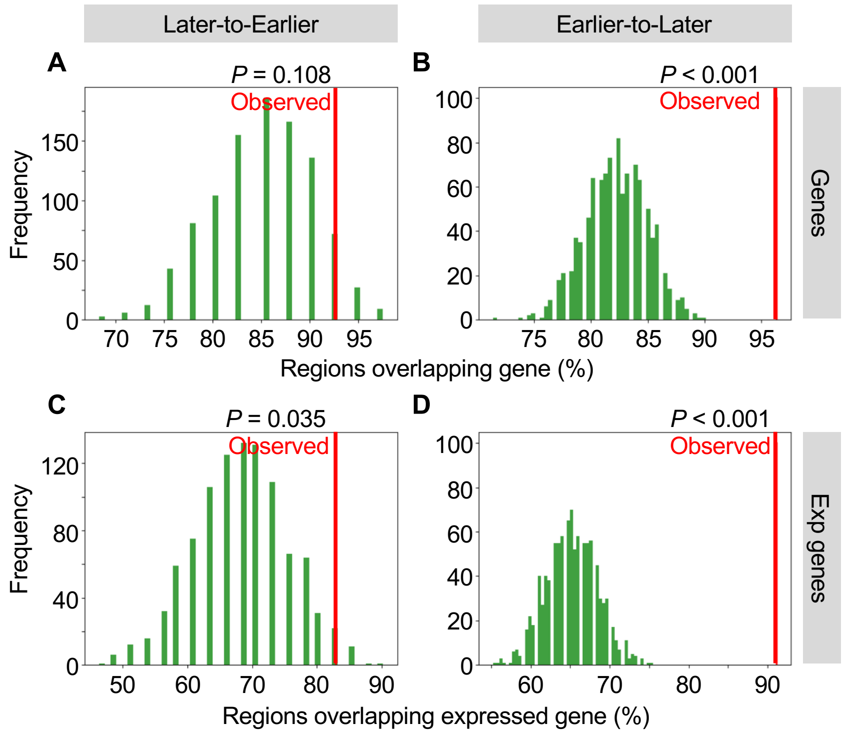

Later-to-Earlier
Earlier-to-Later
A
B
C
D
P = 0.108
Observed
P < 0.001
Observed
P = 0.035
Observed
P < 0.001
Observed
Frequency
Frequency
Genes
Exp genes
Regions overlapping gene (%)
Regions overlapping expressed gene (%)

S8 Fig.

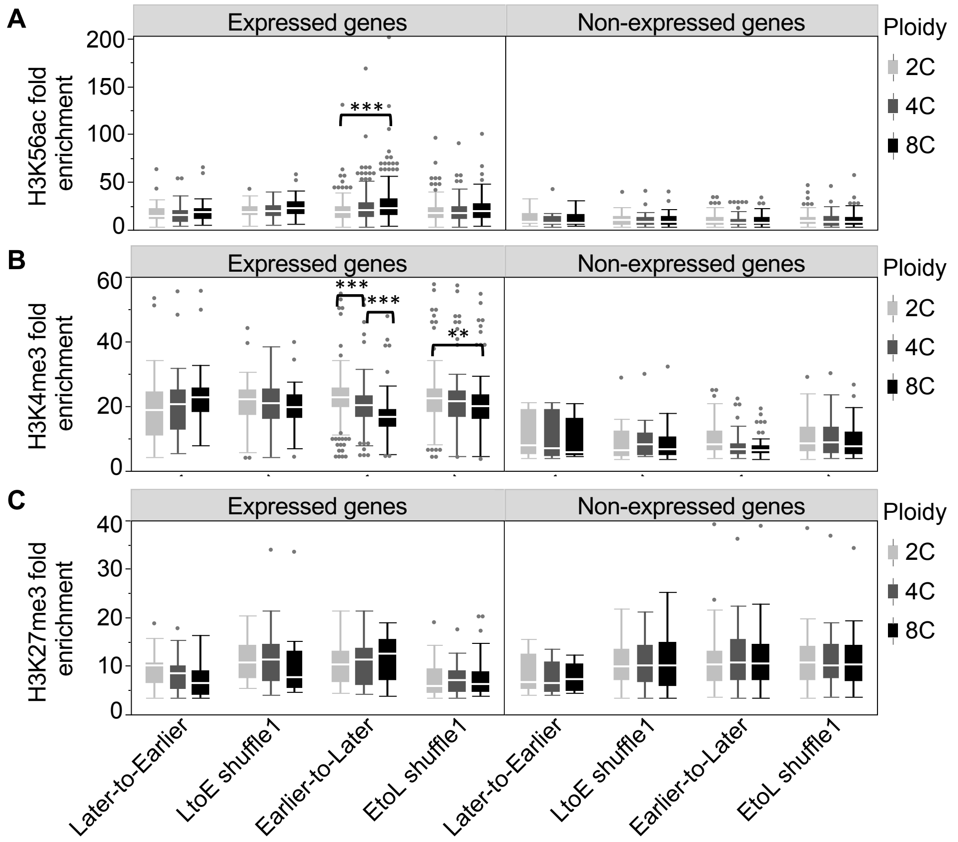

D

| Histone mark | Later-to-Earlier genes | | LtoE shuffle1 genes | | Earlier-to-Later genes | | EtoL shuffle1 genes | |
|---|---|---|---|---|---|---|---|---|
| | Exp (%) | NonExp (%) | Exp (%) | NonExp (%) | Exp (%) | NonExp (%) | Exp (%) | NonExp (%) |
| **H3K56ac** | | | | | | | | |
| 2C | 46 (88.5) | 14 (29.2) | 59 (86.8) | 29 (40.8) | 256 (87.7) | 78 (26.8) | 243 (88.4) | 79 (29.8) |
| 4C | 46 (88.5) | 17 (35.4) | 59 (86.8) | 31 (43.7) | 261 (89.4) | 90 (30.9) | 245 (89.1) | 95 (35.8) |
| 8C | 46 (88.5) | 17 (35.4) | 58 (85.3) | 29 (40.8) | 256 (87.7) | 77 (26.5) | 239 (86.9) | 88 (33.2) |
| 8C/2C Ratio | 1.00 | 1.21 | 0.98 | 1.00 | 1.00 | 0.99 | 0.98 | 1.11 |
| **H3K4me3** | | | | | | | | |
| 2C | 45 (86.5) | 8 (16.7) | 57 (83.8) | 22 (31.0) | 244 (83.6) | 35 (12.0) | 230 (83.6) | 51 (19.2) |
| 4C | 46 (88.5) | 7 (14.6) | 58 (85.3) | 14 (19.7) | 242 (82.9) | 34 (11.7) | 231 (84.0) | 44 (16.6) |
| 8C | 46 (88.5) | 12 (25.0) | 58 (85.3) | 22 (31.0) | 243 (83.2) | 40 (13.7) | 233 (84.7) | 59 (22.3) |
| 8C/2C Ratio | 1.02 | 1.50 | 1.02 | 1.00 | 1.00 | 1.14 | 1.01 | 1.16 |
| **H3K27me3** | | | | | | | | |
| 2C | 21 (40.4) | 14 (29.2) | 13 (19.1) | 28 (39.4) | 20 (6.8) | 82 (28.2) | 35 (12.7) | 87 (32.8) |
| 4C | 19 (36.5) | 14 (29.2) | 13 (19.1) | 27 (38.0) | 21 (7.2) | 80 (27.5) | 31 (11.3) | 84 (31.7) |
| 8C | 17 (32.7) | 14 (29.2) | 13 (19.1) | 28 (39.4) | 22 (7.5) | 86 (29.6) | 33 (12.0) | 89 (33.6) |
| 8C/2C Ratio | 0.81 | 1.00 | 1.00 | 1.00 | 1.10 | 1.05 | 0.94 | 1.02 |
| **Total gene no.** | 52 | 48 | 68 | 71 | 292 | 291 | 275 | 265 |

S9 Fig.

| | | | | | | | | RAT | | | | Random | | | |
|---|---|---|---|---|---|---|---|---|---|---|---|---|---|---|---|
| Gene ontology information | | | | Heatmap | | | | (1) Later-to-Earlier | | (2) Earlier-to-Later | | (3) LtoE shuffle1 | | (4) EtoL shuffle1 | |
| No | GO Term | Onto | Description | 1 | 2 | 3 | 4 | FDR | Num | FDR | Num | FDR | Num | FDR | Num |
| 1 | GO:0010467 | P | gene expression | | | | | --- | --- | 2.5e-10 | 152 | --- | --- | --- | --- |
| 2 | GO:0006139 | P | nucleobase-containing compound metabolic process | | | | | --- | --- | 3.6e-07 | 174 | --- | --- | --- | --- |
| 3 | GO:0006807 | P | nitrogen compound metabolic process | | | | | --- | --- | 6.1e-05 | 195 | --- | --- | --- | --- |
| 4 | GO:0044260 | P | cellular macromolecule metabolic process | | | | | --- | --- | 6.1e-05 | 217 | --- | --- | --- | --- |
| 5 | GO:0043170 | P | macromolecule metabolic process | | | | | --- | --- | 6.1e-05 | 229 | --- | --- | --- | --- |
| 6 | GO:0006412 | P | translation | | | | | --- | --- | 6.1e-05 | 41 | --- | --- | --- | --- |
| 7 | GO:0007049 | P | cell cycle | | | | | --- | --- | 0.00028 | 39 | --- | --- | --- | --- |
| 8 | GO:0043412 | P | macromolecule modification | | | | | --- | --- | 0.00047 | 107 | --- | --- | --- | --- |
| 9 | GO:0016043 | P | cellular component organization | | | | | --- | --- | 0.0083 | 128 | --- | --- | --- | --- |
| 10 | GO:0003723 | F | RNA binding | | | | | --- | --- | 1.1e-12 | 63 | --- | --- | --- | --- |
| 11 | GO:0003676 | F | nucleic acid binding | | | | | --- | --- | 4.6e-08 | 113 | --- | --- | --- | --- |
| 12 | GO:0005198 | F | structural molecule activity | | | | | --- | --- | 3.8e-05 | 26 | --- | --- | --- | --- |
| 13 | GO:0004518 | F | nuclease activity | | | | | --- | --- | 0.0074 | 20 | --- | --- | --- | --- |
| 14 | GO:0005730 | C | nucleolus | | | | | --- | --- | 1.9e-16 | 46 | --- | --- | --- | --- |
| 15 | GO:0044428 | C | nuclear part | | | | | --- | --- | 1.4e-15 | 75 | --- | --- | --- | --- |
| 16 | GO:0031974 | C | membrane-enclosed lumen | | | | | --- | --- | 8.4e-15 | 69 | --- | --- | --- | --- |
| 17 | GO:0043233 | C | organelle lumen | | | | | --- | --- | 8.4e-15 | 69 | --- | --- | --- | --- |
| 18 | GO:0070013 | C | intracellular organelle lumen | | | | | --- | --- | 8.4e-15 | 69 | --- | --- | --- | --- |
| 19 | GO:0031981 | C | nuclear lumen | | | | | --- | --- | 9.1e-14 | 62 | --- | --- | --- | --- |
| 20 | GO:0044424 | C | intracellular part | | | | | --- | --- | 1.3e-12 | 242 | 0.016 | 57 | 0.013 | 208 |
| 21 | GO:0043232 | C | intracellular non-membrane-bounded organelle | | | | | --- | --- | 6.9e-12 | 80 | --- | --- | --- | --- |
| 22 | GO:0043228 | C | non-membrane-bounded organelle | | | | | --- | --- | 6.9e-12 | 80 | --- | --- | --- | --- |
| 23 | GO:0030529 | C | intracellular ribonucleoprotein complex | | | | | --- | --- | 9.1e-12 | 54 | --- | --- | --- | --- |
| 24 | GO:0005622 | C | intracellular | | | | | --- | --- | 4.2e-11 | 243 | 0.034 | 57 | 0.0044 | 216 |
| 25 | GO:0043229 | C | intracellular organelle | | | | | --- | --- | 2.4e-10 | 219 | --- | --- | 0.034 | 184 |
| 26 | GO:0043226 | C | organelle | | | | | --- | --- | 3.4e-10 | 219 | --- | --- | 0.037 | 184 |
| 27 | GO:0043227 | C | membrane-bounded organelle | | | | | --- | --- | 4e-10 | 211 | --- | --- | 0.039 | 175 |
| 28 | GO:0044422 | C | organelle part | | | | | --- | --- | 4.1e-10 | 127 | --- | --- | 0.028 | 96 |
| 29 | GO:0043231 | C | intracellular membrane-bounded organelle | | | | | --- | --- | 5.8e-10 | 210 | --- | --- | 0.039 | 175 |
| 30 | GO:0032991 | C | macromolecular complex | | | | | --- | --- | 1.1e-09 | 103 | --- | --- | 0.039 | 74 |
| 31 | GO:0044446 | C | intracellular organelle part | | | | | --- | --- | 1.5e-09 | 124 | --- | --- | 0.02 | 96 |
| 32 | GO:0044464 | C | cell part | | | | | --- | --- | 1.6e-09 | 253 | 0.0031 | 63 | 0.013 | 227 |
| 33 | GO:0005623 | C | cell | | | | | --- | --- | 2.7e-09 | 253 | 0.0031 | 63 | 0.013 | 228 |
| 34 | GO:0005829 | C | cytosol | | | | | --- | --- | 1.4e-08 | 68 | 0.024 | 17 | --- | --- |
| 35 | GO:0005634 | C | nucleus | | | | | --- | --- | 5e-08 | 135 | --- | --- | --- | --- |
| 36 | GO:0005737 | C | cytoplasm | | | | | --- | --- | 6.2e-08 | 185 | 0.0048 | 48 | 0.013 | 161 |
| 37 | GO:0044444 | C | cytoplasmic part | | | | | --- | --- | 1.5e-07 | 162 | --- | --- | --- | --- |
| 38 | GO:0005654 | C | nucleoplasm | | | | | --- | --- | 6.3e-06 | 28 | --- | --- | --- | --- |
| 39 | GO:0005635 | C | nuclear envelope | | | | | --- | --- | 0.00015 | 13 | --- | --- | --- | --- |
| 40 | GO:0031967 | C | organelle envelope | | | | | --- | --- | 0.00022 | 33 | --- | --- | --- | --- |
| 41 | GO:0005739 | C | mitochondrion | | | | | --- | --- | 0.00026 | 45 | --- | --- | --- | --- |
| 42 | GO:0031975 | C | envelope | | | | | --- | --- | 0.00033 | 33 | --- | --- | --- | --- |
| 43 | GO:0005840 | C | ribosome | | | | | --- | --- | 0.00068 | 24 | --- | --- | --- | --- |
| 44 | GO:0009536 | C | plastid | | | | | --- | --- | 0.0073 | 60 | --- | --- | --- | --- |

Significance heatmap:

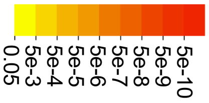

S10 Fig.

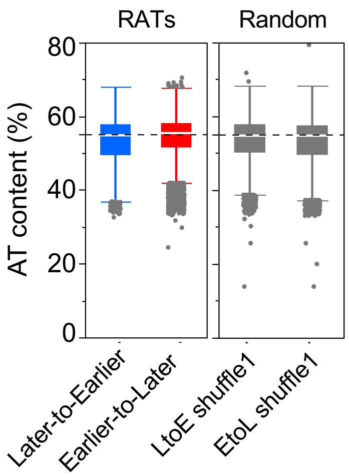
RATs
Random
80
60
40
20
0
AT content (%)
Later-to-Earlier
Earlier-to-Later
LtoE shuffle1
EtoL shuffle1

S11 Fig.

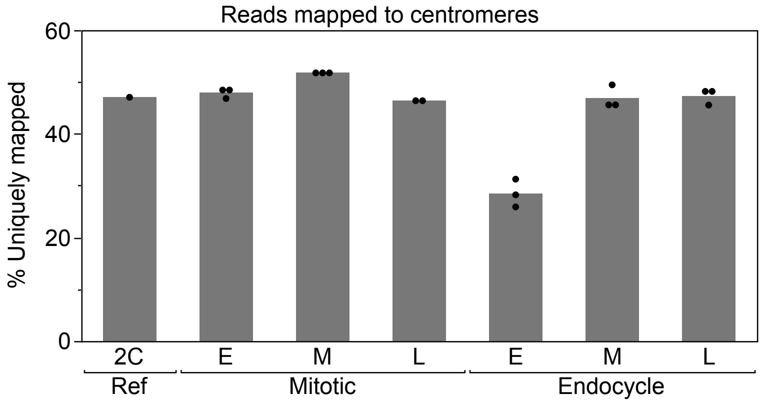

Reads mapped to centromeres
% Uniquely mapped
60
40
20
0
2C
E
M
L
E
M
L
Ref
Mitotic
Endocycle

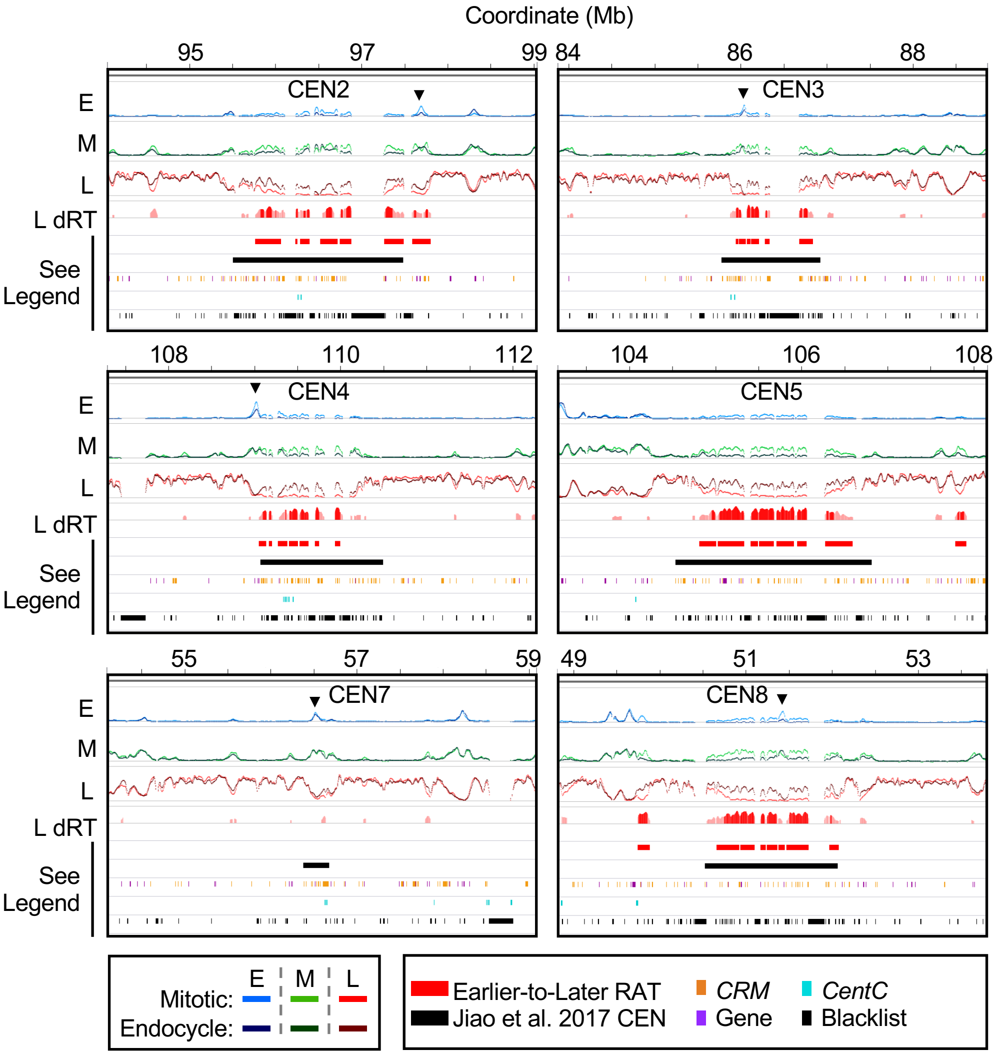
Coordinate (Mb)
95
97
99
84
86
88
CEN2
CEN3
108
110
112
104
106
108
CEN4
CEN5
55
57
59
49
51
53
CEN7
CEN8
E
M
L
L dRT
See Legend
Mitotic:
Endocycle:
E
M
L
Earlier-to-Later RAT
Jiao et al. 2017 CEN
CRM
Gene
CentC
Blacklist

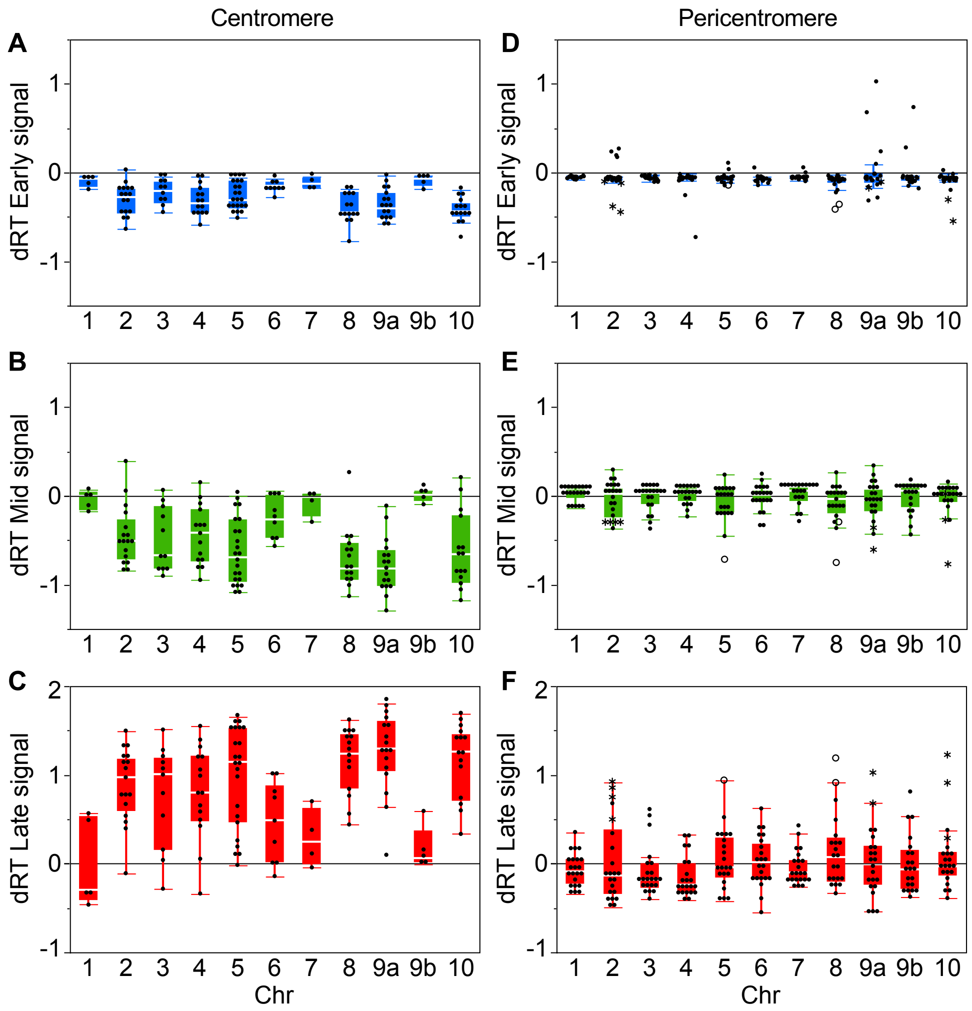
Centromere
Pericentromere
A
B
C
D
E
F
dRT Early signal
dRT Mid signal
dRT Late signal
1
0
-1
2
1 2 3 4 5 6 7 8 9a 9b 10
Chr

S14 Fig.

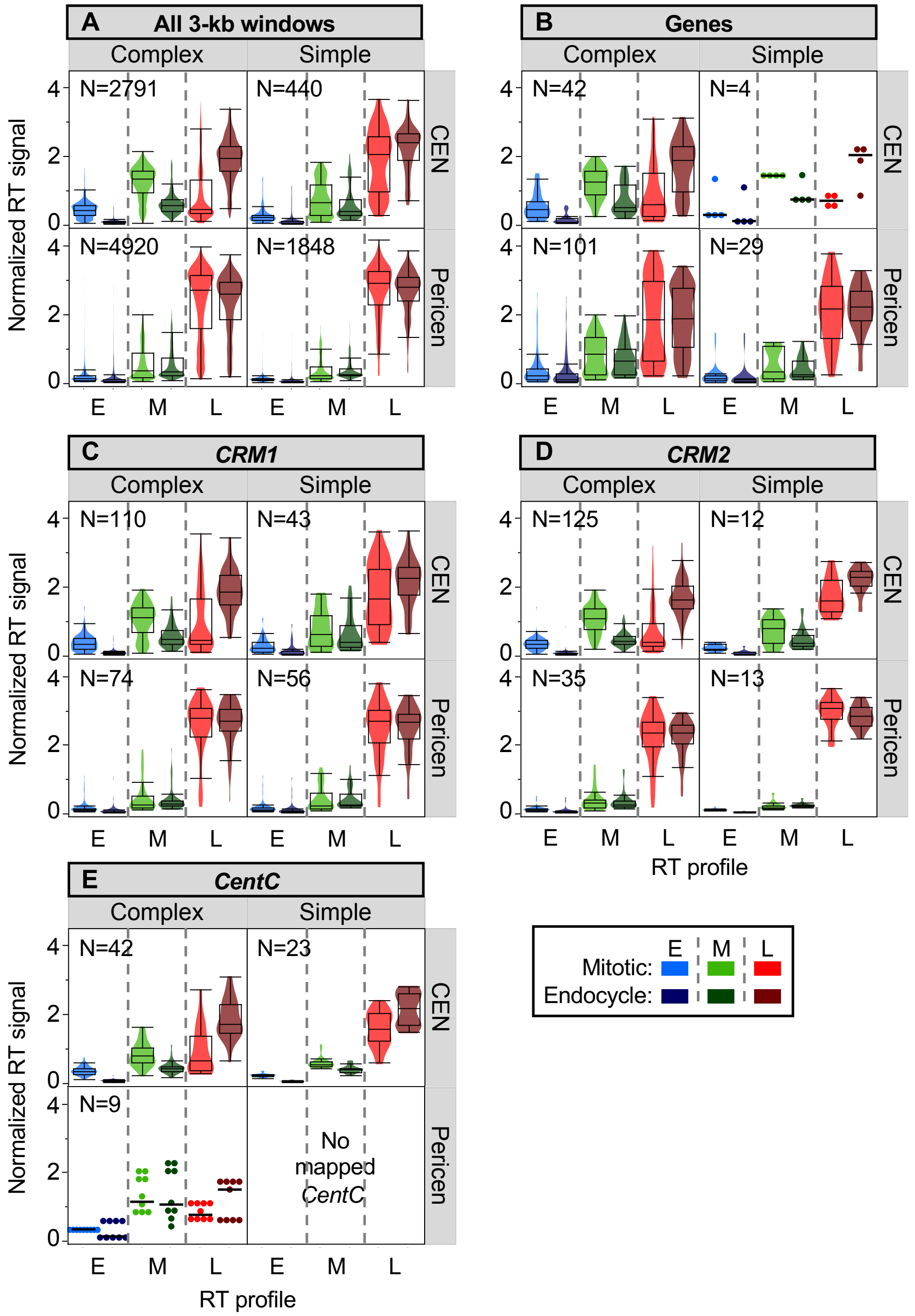
A All 3-kb windows
B Genes
C CRM1
D CRM2
E CentC
Complex
Simple
CEN
Pericen
Normalized RT signal
RT profile
E
M
L
N=2791
N=440
N=4920
N=1848
N=42
N=4
N=101
N=29
N=110
N=43
N=74
N=56
N=125
N=12
N=35
N=13
N=42
N=23
N=9
No mapped CentC
Mitotic:
Endocycle:

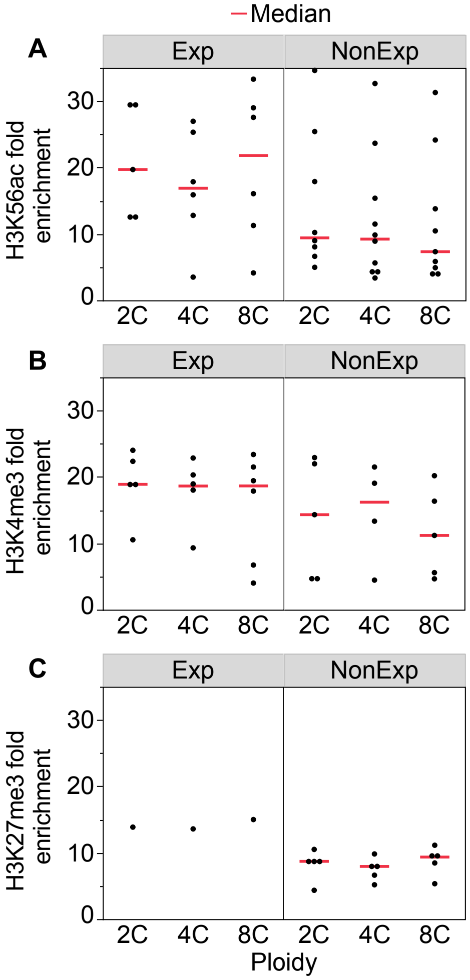

D

| Histone mark | Gene total | Exp | NonExp |
|---|---|---|---|
| **H3K56ac** | | | |
| 2C | 13 | 5 | 8 |
| 4C | 16 | 6 | 10 |
| 8C | 15 | 6 | 9 |
| **H3K4me3** | | | |
| 2C | 10 | 5 | 5 |
| 4C | 9 | 5 | 4 |
| 8C | 11 | 6 | 5 |
| **H3K27me3** | | | |
| 2C | 6 | 1 | 5 |
| 4C | 6 | 1 | 5 |
| 8C | 6 | 1 | 5 |
| **Complex CENs** | 45 | 12 | 33 |

S16 Fig.

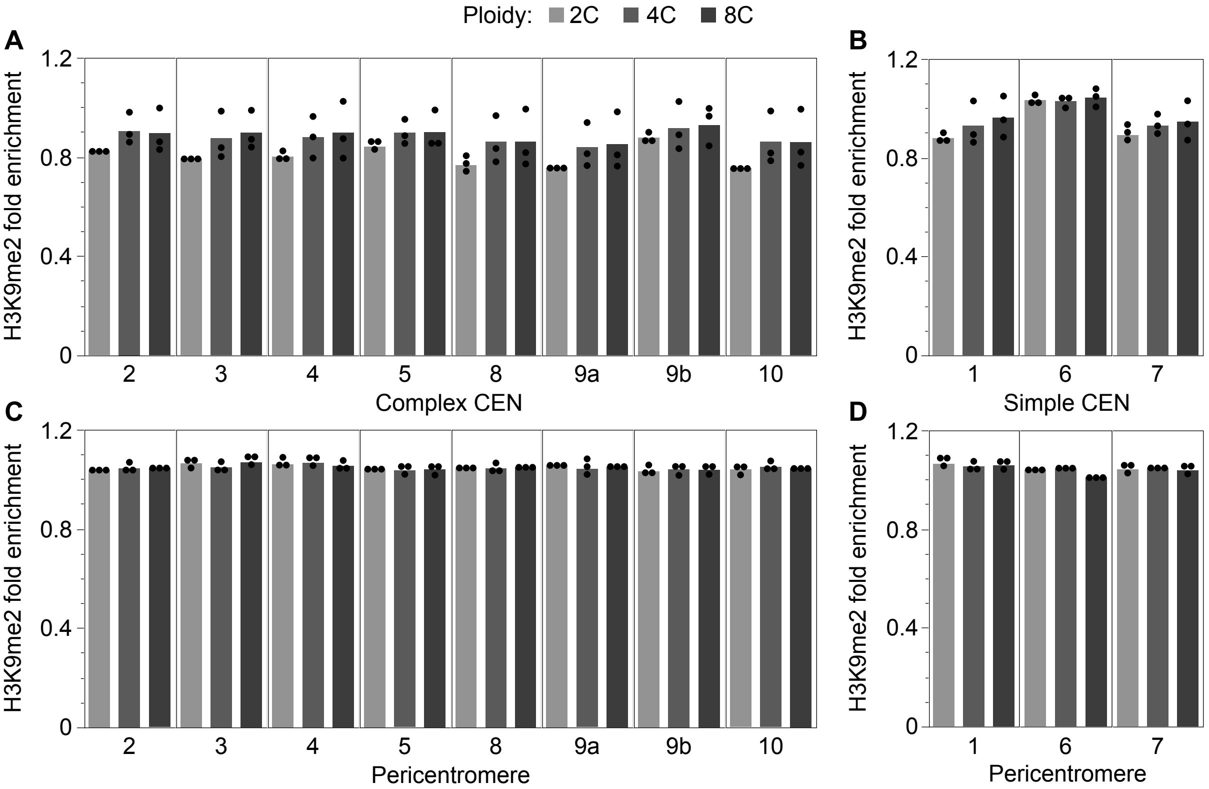
Ploidy: 2C 4C 8C
A
B
C
D
H3K9me2 fold enrichment
1.2
0.8
0.4
0
2
3
4
5
8
9a
9b
10
Complex CEN
1
6
7
Simple CEN
Pericentromere
Pericentromere

S17 Fig.

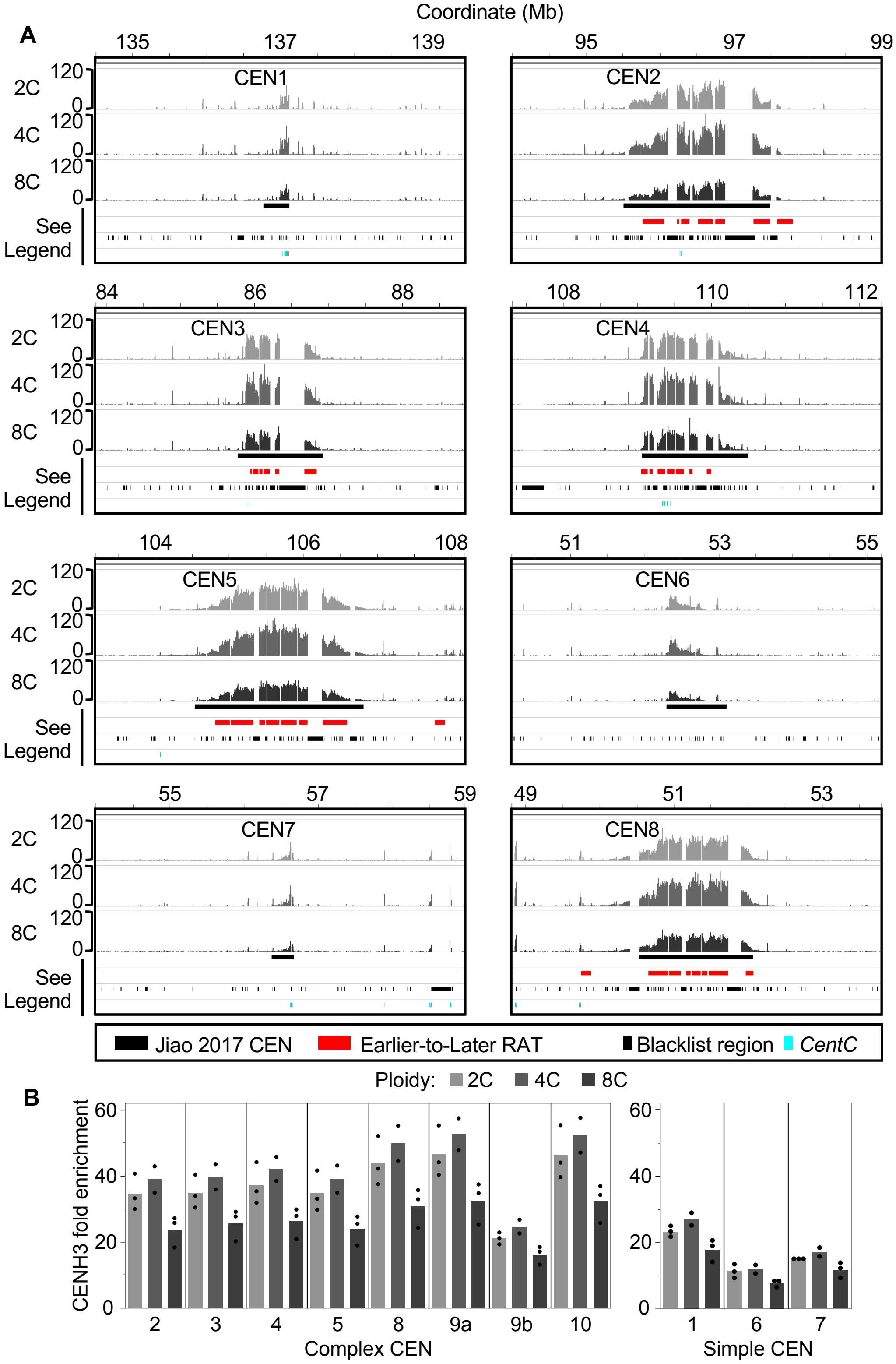
Coordinate (Mb)
A
135
137
139
95
97
99
CEN1
CEN2
120
0
2C
4C
8C
See
Legend
84
86
88
108
110
112
CEN3
CEN4
104
106
108
51
53
55
CEN5
CEN6
55
57
59
49
51
53
CEN7
CEN8
Jiao 2017 CEN
Earlier-to-Later RAT
Blacklist region
CentC
B
Ploidy: 2C 4C 8C
CENH3 fold enrichment
60
40
20
0
2
3
4
5
8
9a
9b
10
Complex CEN
1
6
7
Simple CEN

S18 Fig.

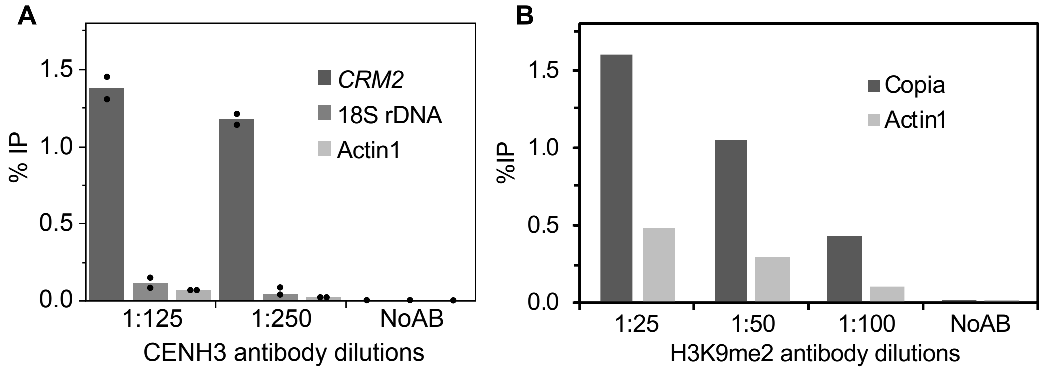
A
1.5
1.0
0.5
0.0
% IP
CRM2
18S rDNA
Actin1
1:125
1:250
NoAB
CENH3 antibody dilutions
B
1.5
1.0
0.5
0.0
%IP
Copia
Actin1
1:25
1:50
1:100
NoAB
H3K9me2 antibody dilutions
