## Supplemental Methods and Tables for "Comparing DNA replication programs reveals large timing shifts at centromeres of endocycling cells in maize roots"

### **SUPPORTING INFORMATION**

**S1 Text. Supplemental Methods.**

#### **Copy number analysis by deep sequencing**

Sequencing data from the CENH3 ChIP-seq input samples (see main text Methods for ChIP-seq data analysis) from nuclei with 2C, 4C and 8C DNA contents were used to look for over- or under-replication by deep sequencing, following the protocol of Yarosh et al. [1]. To gain a better representation of repeat regions in the genome, reads that could not be uniquely mapped to a single location were included in this analysis, but we retained only the primary alignment location for each read pair. After alignment, biological replicate data were scaled to 1× genome coverage, binned in 5-kb static windows and merged by taking the mean of the replicates in each 5-kb window. Ratios were made between 8C and 4C or 4C and 2C data in each 5-kb window. Regions of over- or under-replication were defined as having normalized read frequency ratios that deviated from the mean by two standard deviations in at least two consecutive 5-kb windows, however no genomic windows met this criterion.

**Repeat sequence copy number analysis**

Consensus sequences were acquired for maize repeat classes, including *CentC*, *TR-1*, *knob180*, *CRM* families 1–4 (courtesy of J. Gent and R.K. Dawe) and 5S and 45S rDNA (courtesy of T. Wolfgruber and G. Presting). The abundance of these repeat sequences in the 2C, 4C and 8C input sequence reads was calculated using BLAST software and a non-stringent E value (parameter *-e* 1e-8) as described in Wear et al. [2].

#### **Antibody validations**

ChIP-qPCR experiments were used to validate the specificity and optimal titration of anti-H3K9me2 and *Zea mays* anti-CENH3 (gift from R.K. Dawe laboratory) antibodies using fixed maize root material (S18 Fig). Antibody dilutions for H3K27me3, H3K4me3 and H3K56ac were determined previously [2]. ChIP procedures were carried out as described in main text Methods, except whole fixed root material (without nuclei isolation or flow sorting) was used. PCR reactions were carried out in triplicate using Power SYBR Green PCR Master Mix (Applied Biosystems), and included no antibody and no template negative controls. The *CRM2* primer sequences (courtesy of J. Gent) were CTAGGACTTGCTGGCTTCTATC (JIG-261) and TCCCTTCTTCGTAAGCTCATTC (JIG-262). The 18S rDNA primers were ACTGTGAAACTGCGAATGGC and GAAGTCGGGGTTTGTTGCAC. The Copia reverse transcriptase and Actin1 UTR primers were from Haring et al. [3]. The percentage of input (%IP) [3] was calculated for various antibody dilutions and a no antibody control.

#### **ChIP-seq data peak calling and analysis**

H3K56ac, H3K4me3 and H3K27me3 peaks of enrichment were called from ChIP-seq data generated from nuclei with 2C, 4C or 8C DNA content using MACS v2.1.0 *callpeak* [4] with parameters “–nomodel–nolambda–broad” and a q-value threshold of 0.01. Peaks were called from merged and individual biological replicate files, and only “replicated” peaks observed in all three biological replicates were retained. The intersection of 2C, 4C and 8C peaks with genes in non-CEN RATs or in centromeres was determined using intersectBED in the BEDtools suite (parameters *-wa -wb*) [5]. If multiple peaks were found on a single gene, their peak fold enrichment values from the MACS2 output were summed using mergeBED (parameter -*o* sum). We applied a non-parametric Kruskal-Wallis analysis of variance by rank test for significant differences in histone modification fold enrichment values between ploidy levels in SAS JMP Pro v14 (SAS Institute Inc.). If a significant *P* value of < 0.05 was obtained, a Steel-Dwass-Critchlow-Fligner post-hoc test was carried out between each pairwise comparison.

#### **RAT gene ontology analysis**

“Expressed” genes were defined as having a FPKM (fragments per kilobase of transcript per million mapped reads) level of ≥ 1 in at least one of two RNA-seq datasets previously generated in our lab from maize seedling roots of the same age and under the same growth conditions as used for the Repli-seq and ChIP-seq experiments. The RNA-seq datasets were derived from whole root sections of 0–1 mm [2] and 0–5 mm, and were prepared and analyzed as described in Wear et al. [2], except that reads were mapped to B73 RefGen_v4 Zm00001d.2 gene annotations. Lists of expressed genes found in Earlier-to-Later and Later-to-Earlier non-CEN RATs were used for gene ontology (GO) analysis with the agriGO v2.0 cross-comparison singular enrichment analysis (SEACOMPARE) tool and default parameters [6], except that we used the Plant GO slim subset developed by The Arabidopsis Information Resource (TAIR; https://www.arabidopsis.org) to provide a broad representation of GO categories. The RefGen_v4 gene GO term reference set was assigned by maize-GAMER [7].

### **SUPPLEMENTAL TABLES**

**S1 Table. Replication timing signal differences and thresholds.**

| **S-phase fraction** | **Average across chromosomes** | | | **10% threshold** | **25% threshold** |
| --- | --- | --- | --- | --- | --- |
|  | **Negative difference max** | **Positive difference max** | **Total difference range** |  |  |
| **Early** | -2.6 | 1.9 | 4.5 | -- | -- |
| **Late** | -1.6 | 1.9 | 3.5 | -- | -- |
| **Average** | -2.1 | 1.9 | 4.0 | 0.4 | 1.0 |

**S2 Table. Summary statistics of preliminary RAT calling steps.**

| **RT change** | **Stringency threshold** | **Region count** | **Min size (kb)** | **Median size (kb)** | **Max size (kb)** | **Sum (kb)** | **Genome coverage (%)** |
| --- | --- | --- | --- | --- | --- | --- | --- |
| **Earlier-to-Later** | ≥ 10% | 2626 | 3 | 39 | 1110 | 144,996 | 6.9 |
|  | ≥ 25% | 284 | 3 | 27 | 543 | 12,468 | 0.6 |
|  | **Final RATs** | **233** | **24** | **135** | **1110** | **34,575** | **1.6** |
| **Later-to-Earlier** | ≥ 10% | 2180 | 3 | 30 | 357 | 87,936 | 4.2 |
|  | ≥ 25% | 49 | 3 | 24 | 102 | 1,494 | 0.1 |
|  | **Final RATs** | **41** | **60** | **141** | **315** | **6,291** | **0.3** |

**S3 Table. Gene summary in non-centromeric RATs.**

|  | **Region count** | **Genome coverage (%)** | **RATs with gene (%)**^a^ | **RATs with expressed gene (%)**^a^ | **Gene count** | | |
| --- | --- | --- | --- | --- | --- | --- | --- |
|  |  |  |  |  | **Total** | **Expressed^b^**  **(% of total)** | **Mean No.**  **per region^c^** |
| **Later-to-Earlier** | 41 | 0.3 | 38 (92.7) | 34 (82.9) | 100 | 52 (52.0) | 2.6 |
| **Earlier-to-Later** | 192 | 1.3 | 185 (96.4) * | 175 (91.1) * | 582 | 292 (50.2) | 3.2 |
| ***Total*** | 233 | 1.6 | 223 (95.7) | 209 (89.7) | 682 | 344 (50.4) | 3.1 |
| Notes:  ^a^ Asterisks indicate values that were significantly greater than random expectation (P value ≤ 0.001) from permutation analysis. See S7 Fig.  ^b^ Expressed genes were defined as those having an FPKM ≥ 1 in at least one of two root specific RNA-seq  datasets (see S1 Text for details).  ^c^ The mean count of genes per region (including all genes). | | | | | | | |

**S4 Table. Permutation analysis results for gene and TE coverage in non-CEN RATs.**

|  | **Later-to-Earlier RAT** | | **Earlier-to-Later RAT** | | **Whole genome** |
| --- | --- | --- | --- | --- | --- |
| **Genes** | **% Coverage^a^** | ***P* value** | **% Coverage^a^** | ***P* value** | **% Coverage^a^** |
| All | 4.97 | NEG 0.009 | 10.32 | **≤ 0.001** | 7.80 |
| Expressed (FPKM > 1) | 3.52 | NEG 0.021 | 7.94 | **≤ 0.001** | 5.51 |
| **TE Superfamily**  **/ Common name** | **% Coverage^a^** | ***P* value** | **% Coverage^a^** | ***P* value** | **% Coverage^a^** |
| RLX / Unknown LTR | 8.45 | 0.286 | 7.43 | 0.684 | 7.90 |
| RLG / Gypsy | 41.02 | 0.019 | 31.83 | **NEG**  **≤ 0.001** | 36.38 |
| RLC / Copia | 15.07 | NEG 0.023 | 20.12 | 0.095 | 19.02 |
| RIT / LINE RTE | 0.00 | NA^b^ | 0.03 | 0.279 | 0.02 |
| RIL / LINE L1 | 0.08 | 0.531 | 0.11 | 0.414 | 0.10 |
| RST / SINE | 0.02 | 0.267 | 0.02 | 0.207 | 0.01 |
| DTX / Unknown TIR | 0.64 | 0.817 | 0.77 | 0.670 | 0.82 |
| DTT / Tc1-Mariner | 1.18 | 0.236 | 1.26 | 0.008 | 1.09 |
| DTM / Mutator | 0.01 | 0.405 | 0.01 | NEG 0.069 | 0.03 |
| DTH / PIF-Harbinger | 0.93 | 0.644 | 0.94 | 0.723 | 0.98 |
| DTC / CACTA | 0.54 | 0.268 | 0.38 | 0.620 | 0.42 |
| DTA / hAT | 0.10 | 0.888 | 0.20 | 0.206 | 0.17 |
| DHH / Helitron | 2.86 | NEG 0.044 | 4.49 | 0.059 | 4.01 |
| TE total | 70.90 | 0.338 | 67.58 | NEG 0.009 | 70.97 |
| Notes:  ^a^ Cumulative percent coverage.  ^b^ No overlap was found, so a *P* value was not calculated. | | | | | |

**S5 Table. Cumulative RAT coverage in centromeres.**

| **CEN** | **CEN size total (kb)^a^** | **Included 3-kb window count** | **CEN excluding blacklist (kb)^b^** | **Total RAT**  **coverage (kb)^c^** | **RAT count^d^** |
| --- | --- | --- | --- | --- | --- |
| 1 | 350 | 102 | 305 | 0 | 0 |
| 2 | 1980 | 379 | 1137 | 1098 | 7 |
| 3 | 1150 | 196 | 587 | 405 | 6 |
| 4 | 1430 | 285 | 854 | 495 | 7 |
| 5 | 2280 | 548 | 1644 | 1284 | 7 |
| 6 | 800 | 241 | 722 | 0 | 0 |
| 7 | 300 | 92 | 276 | 0 | 0 |
| 8 | 1540 | 361 | 1082 | 939 | 7 |
| 9a | 1650 | 494 | 1482 | 1518 | 4 |
| 9b | 400 | 93 | 280 | 0 | 0 |
| 10 | 1390 | 432 | 1297 | 1338 | 3 |
| Notes:  ^a^ Centromere size reported in [8], which includes unmappable regions of known size.  ^b^ Centromere size, excluding blacklist regions (unmappable and multi-mapping).  ^c^ The combined coverage of RATs called in each centromere, not including blacklist regions. Note, in some centromeres, called RATs extends past the previously reported CEN boundary.  ^d^ The number of individual RATs called in each centromere. | | | | | |

**S6 Table. Compensated timing shifts in complex centromeres and corresponding pericentromeres.**

|  | **Total 3-kb**  **window count** | **Earlier-to-Later shift** | | **Later-to-Earlier shift** | |
| --- | --- | --- | --- | --- | --- |
|  |  | **Windows compensated**  **at** ≥ **10%** | **% of total** | **Windows compensated**  **at** ≥ **10%** | **% of total** |
| **Centromeres** | 2,791 | 2,384 | 85.4 | 0 | 0.0 |
| **Pericentromeres** | 4,757^a^ | 372 | 7.8 | 77 | 1.6 |
| Notes:  ^a^ Windows that were clearly an extension of a centromere RAT, but were outside the published centromere boundary were excluded from the pericentromere counts. | | | | | |

**S7 Table. CENH3 average fold enrichment relative to DNA content in centromeres.**

| **CEN** | **CENH3 average fold enrichment (± S. D.)** | | | **Ratio values** | |
| --- | --- | --- | --- | --- | --- |
|  | **2C** | **4C** | **8C** | **4C/2C** | **8C/4C** |
| 1 | 23.1 ± 1.5 | 27.0 ± 2.7 | 17.7 ± 3.2 | 1.17 | 0.73 |
| 2 | 36.2 ± 5.8 | 40.7 ± 5.9 | 24.6 ± 4.8 | 1.11 | 0.68 |
| 3 | 36.2 ± 5.3 | 41.2 ± 5.7 | 26.5 ± 4.9 | 1.13 | 0.72 |
| 4 | 39.5 ± 6.7 | 44.8 ± 5.5 | 27.9 ± 5.0 | 1.12 | 0.69 |
| 5 | 35.6 ± 6.3 | 39.9 ± 5.9 | 24.6 ± 4.6 | 1.11 | 0.69 |
| 6 | 11.4 ± 2.1 | 12.0 ± 1.9 | 7.8 ± 1.1 | 1.06 | 0.71 |
| 7 | 15.2 ± 0.6 | 17.3 ± 1.9 | 11.7 ± 2.1 | 1.13 | 0.75 |
| 8 | 45.1 ± 7.7 | 51.2 ± 7.7 | 31.7 ± 6.1 | 1.13 | 0.70 |
| 9a | 46.9 ± 7.8 | 53.0 ± 6.8 | 32.7 ± 6.4 | 1.11 | 0.69 |
| 9b | 21.3 ± 1.9 | 24.9 ± 3.0 | 16.3 ± 2.7 | 1.17 | 0.72 |
| 10 | 46.7 ± 8.2 | 52.8 ± 7.5 | 32.5 ± 5.9 | 1.11 | 0.69 |
| **Theoretical values:** | | | | | |
| **Proportional redeposition** | | | | 1.0 | 1.0 |
| **No redeposition** | | | | 0.5 | 0.5 |
